# OHP2 is not required for *psbA* translation in *Chlamydomonas*

**DOI:** 10.1101/2022.08.29.505686

**Authors:** Fei Wang, Korbinian Dischinger, Lisa Désirée Westrich, Irene Meindl, Felix Egidi, Raphael Trösch, Frederik Sommer, Xenie Johnson, Michael Schroda, Joerg Nickelsen, Felix Willmund, Olivier Vallon, Alexandra-Viola Bohne

**Author notes:** Present address: CEA, BIAM, Aix-Marseille Université, Saint-Paul-lez-Durance, F-13108, France. Contact information: Olivier Vallon and Alexandra-Viola Bohne.

## Abstract

In land plants and cyanobacteria, co-translational association of chlorophyll (Chl) to the nascent D1 polypeptide, a reaction center protein of photosystem II (PSII), requires a Chl binding complex consisting of a short-chain dehydrogenase (HCF244/Ycf39) and One-Helix Proteins of the LHC superfamily (OHP1 and OHP2 in chloroplasts). Here, we show that an *ohp2* mutant of the green alga *Chlamydomonas reinhardtii* fails to accumulate core PSII subunits, in particular D1. Extragenic suppressors arise at high frequency, suggesting the existence of another route for Chl association to PSII. The *ohp2* mutant can be complemented by the *Arabidopsis* ortholog. In contrast to land plants, where *psbA* translation is prevented in the absence of OHP2, ribosome profiling experiments show that the *Chlamydomonas* mutant translates the *psbA* transcript over its full length. Pulse labelling suggests that D1 is degraded during or immediately after translation. The translation of other PSII subunits is affected by assembly-controlled translational regulation (the CES process). Proteomics show that HCF244, a translation factor which associates with and is stabilized by OHP2 in land plants, still partly accumulates in the *Chlamydomonas ohp2* mutant, explaining the persistence of *psbA* translation. Several Chl biosynthesis enzymes overaccumulate in the mutant membranes. Partial inactivation of the D1-degrading FtsH protease restores a low level of PSII activity in an *ohp2* background, but not photoautotrophy. Taken together, our data suggest that OHP2 is not required for *psbA*D1 translation in *Chlamydomonas*, but necessary for its stabilization.

## INTRODUCTION

Oxygenic photosynthesis, carried out by cyanobacteria and photosynthetic eukaryotes, is based on the light-driven electron transfer from water to NADPH, involving three complexes located in the thylakoid membrane and operating in series: photosystem II (PSII), cytochrome *b*_6_*f* (Cyt*b_6_f*) and photosystem I (PSI). PSII is a multi-subunit complex, whose core complex is made up of 20-23 subunits: the reaction center proteins D1 (PsbA), D2 (PsbD), Cyt*b*_559_ (subunits PsbE and PsbF) and PsbI are encoded in the chloroplast genome by the *psbA, psbD, psbE/F* and *psbI* genes, as are the core antenna CP47 (PsbB) and CP43 (PsbC) and the additional subunits PsbH, PsbJ, PsbK, PsbL, PsbM, PsbT and PsbZ (reviewed in e.g. Gao et al., 2018). In eukaryotes, nucleus-encoded subunits complete the complex and build its peripheral chlorophyll (Chl) *a/b* containing light-harvesting antenna complexes (LHCs). Chl *a* and its derivative pheophytin (Phe) *a* are essential components of the intra-PSII electron transfer chain that allows the generation of a stable charge separated state after light-capture. The pigments and redox-active cofactors are scaffolded mainly by the homologous D1 and D2 polypeptides, which also provide Tyr residues involved in re-reduction of P680^+^ and residues liganding the manganese (Mn)-cluster where water-splitting occurs.

Given the tight folding of the enzyme and the high chemical reactivity of Chl cation radicals that can form from the light-excited states, it has long been hypothesized that specific mechanisms must coordinate the synthesis of the reaction center proteins, their intra-membrane insertion and their association with pigments. Over the years, the biogenesis of PSII has been dissected into a series of discrete assembly steps (reviewed in Nickelsen and Rengstl, 2013). Precursor D1 (pD1), co-translationally inserted into the membrane, binds to PsbI and specific assembly factors, forming a D1 module that will associate with a D2 module formed between D2, Cyt*b*_559_ and other factors (Knoppová et al., 2022; Komenda et al., 2012). The reaction center (RC) subcomplex will first integrate CP47, then CP43, associated with other small subunits, forming the monomeric PSII core. Dimerization ensues, as well as association with LHCs integrated into the membrane by the CpSRP complex. On the luminal surface of the enzyme, cleavage of the C-terminal extension of pD1 and light-driven assembly of the Mn-containing cluster allow formation of the water-splitting PSII enzyme.

Over the years, numerous proteins have been found to catalyze various steps of this pathway, both during *de novo* assembly and during repair after photoinhibition (Heinz et al., 2016; Lu, 2016). Some act by regulating gene expression, in particular translation of *psbA* which codes for the rapidly-turning over D1. Others act as a chaperone, binding an assembly intermediate until the next step can be completed. In this category, particular attention has been paid to proteins that could mediate the assembly of the cofactors. It is believed that Chl molecules are presented to the acceptor PSII subunits by specific carrier proteins that precisely mediate their insertion at the proper position, at the proper stage of assembly.

In *Arabidopsis*, a complex consisting of two conserved One-Helix Proteins, OHP1 and OHP2, and a protein related to short-chain dehydrogenase/reductases, HCF244, has been found to be essential for Chl integration into PSII, or for protection of the newly synthesized Chl-associated D1 during formation of the RC complex (Chotewutmontri and Barkan, 2020; Chotewutmontri et al., 2020; Hey and Grimm, 2018; Li et al., 2019; Myouga et al., 2018). In combination with OHP1, OHP2 is able to bind Car and Chl, the latter via specific residues of a Chl binding motif (Hey and Grimm, 2020), and the reconstituted heterodimer has unique photoprotective properties (Psencik et al., 2020). The **O**HP1/OHP2/**H**CF244 **C**omplex (which we will hereafter call “OHC”) is homologous to a complex of similar function described in cyanobacteria, where the relative stability of the assembly intermediates in core subunits mutants has allowed a fine dissection of the pathway (Komenda et al., 2012). OHP1 and OHP2 resemble cyanobacterial High Light-Inducible Proteins (HLIPs) encoded by the *hliA-D* genes (Komenda and Sobotka, 2016), in that they all present a single transmembrane domain showing the key residues for binding Chl(Engelken et al., 2010). While OHP1 clearly derives from HLIPs, the origin of OHP2 is less clear (Engelken et al., 2010). HliC and HliD have been found, together with the HCF244 homolog Ycf39, to associate with the D1 module and then to the RC subcomplex after binding of the D2 module (Chidgey et al., 2014; Knoppová et al., 2014). HliD can bind Chl and β-carotene (Staleva et al., 2015). The Chl synthase ChlG which carries an N-terminal HLIP domain co-immunoprecipitates with HliD/Ycf39 and it is believed that the complex can deliver newly synthesized Chl to D1 and/or D2 during or early after their translation (Chidgey et al., 2014; Staleva et al., 2015). For this process, Chl could also be scavenged from other Chl-binding proteins upon their degradation.

The phenotype of land plant mutants has revealed an additional function of the OHC, namely to regulate *psbA* translation. *Arabidopsis ohp1*, *ohp2* and *hcf244* mutants show strongly reduced PSII levels, while only weak or indirect effects are observed on PSI accumulation (Beck et al., 2017; Li et al., 2019; Link et al., 2012; Myouga et al., 2018). In the three mutants, the other two partners of the OHC are undetectable or dramatically reduced, suggesting that formation of the complex is required for their stabilization (Beck et al., 2017; Hey and Grimm, 2020; Li et al., 2019). Interestingly, *ohp1* and *ohp2* mutants from *Arabidopsis* as well as an *hcf244* mutant from maize show a strong reduction (resp. 7-fold, 12-fold and 11-fold) of the ribosome footprint RF-Seq signal (RF-Seq; Chotewutmontri et al., 2020), indicating near complete inhibition of *psbA* translation(RF-Seq; Chotewutmontri et al., 2020). This has led to the hypothesis that the OHC regulates *psbA* translation by sensing the production of D1 properly loaded with Chl: as long as D1 remains bound to the OHC, HCF244 is unable to catalyze *psbA* translation initiation and D1 translation is halted (Chotewutmontri and Barkan, 2020). Such a tight coupling between assembly and translation is reminiscent of a process described as Control by Epistasy of Synthesis (CES) in *Chlamydomonas* (Minai et al., 2006; Wollman et al., 1999). In the PSII CES cascade, unassembled D1, as produced for example in the absence of D2, represses its own translation initiation. It was thus of particular interest to determine whether such an OHC-dependent translational regulation exists in *Chlamydomonas*.

Many other proteins of the PSII assembly pathway are conserved between cyanobacteria, land plants and *Chlamydomonas*, but their detailed working mechanism has only rarely been investigated in the alga (Spaniol et al., 2021). Among those acting at a site close to that of the OHC, another short-chain dehydrogenase/reductase, HCF173, has been found to be necessary for D1 synthesis and to bind the *psbA* 5’ UTR, along with another protein of ill-defined function, SRRP1 (Link et al., 2012; McDermott et al., 2019; Schult et al., 2007; Watkins et al., 2020). ALB3, which belongs to a conserved family of protein integrases also found in bacteria and mitochondria, plays a major role in the insertion of integral thylakoid membrane proteins in general, and serves as a hub for many other factors (reviewed in Plöchinger et al., 2016). In the case of D1, ALB3 appears to interact with factors acting downstream of D1 integration, namely HCF136, but also LPA1 and probably LPA2 and LPA3. HCF136 (and its homolog Ycf48 in cyanobacteria) is involved, along with the chloroplast-encoded PsbN, in the interaction between the D1 and D2 subcomplexes and thus the formation of the RC subcomplex (Komenda et al., 2008; Plücken et al., 2002; Torabi et al., 2014). Further downstream, LPA2 catalyzes a step leading to the formation of stable PSII monomers, probably by facilitating the integration of CP43 into the RC47 subcomplex (Cecchin et al., 2021; Schneider et al., 2014; Spaniol et al., 2021).

Another factor that plays an important role in the early steps of PSII assembly is a rubredoxin-like protein (Calderon et al., 2013). Here, the cyanobacterial RubA, like the accessory factor Ycf48, has been shown to be a component of the initial D1 assembly module (Kiss et al., 2019). The homologous protein RBD1 from *Chlamydomonas* was proposed to participate in the protection of PSII intermediate complexes from photooxidative damage during de novo assembly and repair (García-Cerdán et al., 2019). Moreover, Calderon et al. (2022) propose a model in which RBD1 promotes the proper folding of D1, possibly via delivery or reduction of the non-heme iron during PSII assembly. Besides a function in the early steps of PSII assembly, RBD1 from *Arabidopsis* was additionally proposed to play a role in the translation of the *psbA* mRNA (Che et al., 2022). Chl insertion into nascent D1 is not only required during PSII biogenesis, but also during repair after photoinhibition. The D1 protein is the primary target of photodamage, which includes a reversible component, repaired in the absence of protein translation, and an irreversible component that requires degradation of D1 and *de novo* synthesis (for a review see e.g. Theis and Schroda, 2016). FtsH is a multi-subunit ATP-dependent thylakoid membrane metalloprotease, combining chaperone and peptidase domains. It has been shown to be a major player in the degradation of D1 and is thus essential for repair after photoinhibition (Lindahl et al., 2000; Silva et al., 2003).(van Wijk, 2015)In *Chlamydomonas*, *ftsH1* mutants show light-sensitivity and defects in PSII repair (Malnoë et al., 2014). The mutants accumulate damaged D1 and its cleavage products generated by DEG-family endopeptidases. How D1 is extracted from PSII without its total disassembly, and how degradation is coupled with synthesis of a replacement subunit, is still largely unknown. In this study, we show that a null mutant of OHP2 in *Chlamydomonas* is entirely devoid of PSII. The primary defect is a lack of D1 accumulation which can be fully restored by complementation of the mutant with *Arabidopsis* OHP2. Ribosome profiling and pulse labeling experiments show that this defect is not caused by an arrest of *psbA* translation, but by a reduced stability of the nascent D1 protein. Targeted proteomics show that HCF244 accumulates to ~25% of WT in the *ohp2* mutant, explaining this partial uncoupling between the association of D1 with Chl and the initiation of translation on *psbA*. In the mutant, D1 degradation appears in part mediated by the FtsH protease. An intriguing specificity of *Chlamydomonas* OHP2 is the high-frequency suppression of the PSII-less phenotype in the mutant.

## RESULTS

### Isolation of a *Chlamydomonas* photosynthetic mutant and identification of the mutated locus

By nuclear transformation of the antibiotic resistance marker *aphVIII*, conferring resistance to paromomycin (Pm), we previously generated a collection of insertional mutants of *Chlamydomonas* (Houille-Vernes et al., 2011). The primary screening of this collection combined analysis of fluorescence induction curves with sorting of photosynthetic versus non-photosynthetic (acetate-requiring) phenotypes to categorize the affected photosynthetic complex. The strain*10.1a* was identified as acetate requiring and deficient in PSII. This strain carries a single insertion of the *aphVIII-*cassette in gene *Cre11.g480000* as revealed by inverse PCR and Southern blotting (Supplemental Figure S1A). In a backcross of *10.1a* to the WT strain WT-S34 (mt*+*), the PSII-deficient phenotype segregated 2:2, indicating a single nuclear mutation, but it was neither linked to Pm resistance nor to the cassette inserted in *Cre11.g480000* (Supplemental Figure S1B, top row). This suggested that a second nuclear mutation causes the observed PSII phenotype.

Strikingly, all of the 47 PSII-deficient progenies analyzed carried the mating type minus (mt-) gene *MID*, and none the mt+ specific *FUS1* (Supplemental Figure S1B, bottom). This indicates linkage to the mating type locus, a ~1 Mb region of recombinational suppression involved in sex determination, residing on chromosome 6 and comprising about 35 genes, many unrelated to sexuality (Ferris et al., 2002). Illumina genome sequencing revealed near the MT locus a single structural variant at position chromosome_6:305949 (v5 genome assembly), i.e. within exon 3 of the *OHP2* gene (*Cre06.g251150*, Figure 1A; Supplemental Figure S2). Analysis of reads flanking this site and their mate reads pointed to the long terminal repeat (LTR) sequence of *TOC1* (Transposon Of Chlamydomonas, Day et al., 1988), indicating a *TOC1* insertion (Supplemental Data S1). Accordingly, Southern blot analysis of the *10.1a* mutant showed integration of a large DNA fragment of ~6 kb into *OHP2* (Figure 1B) and none of the 47 PSII-deficient progenies described above gave rise to an *OHP2*-specific PCR-product (Supplemental Figure S1C). To exclude an impact of the initially identified mutation in *Cre11.g480000* on the mutant phenotype, most analyses were performed with a strain of the backcrossed progeny, which we will term *ohp2*, possessing only the mutation in *OHP2* but not the one in *Cre11.g480000*.

**Figure 1.**
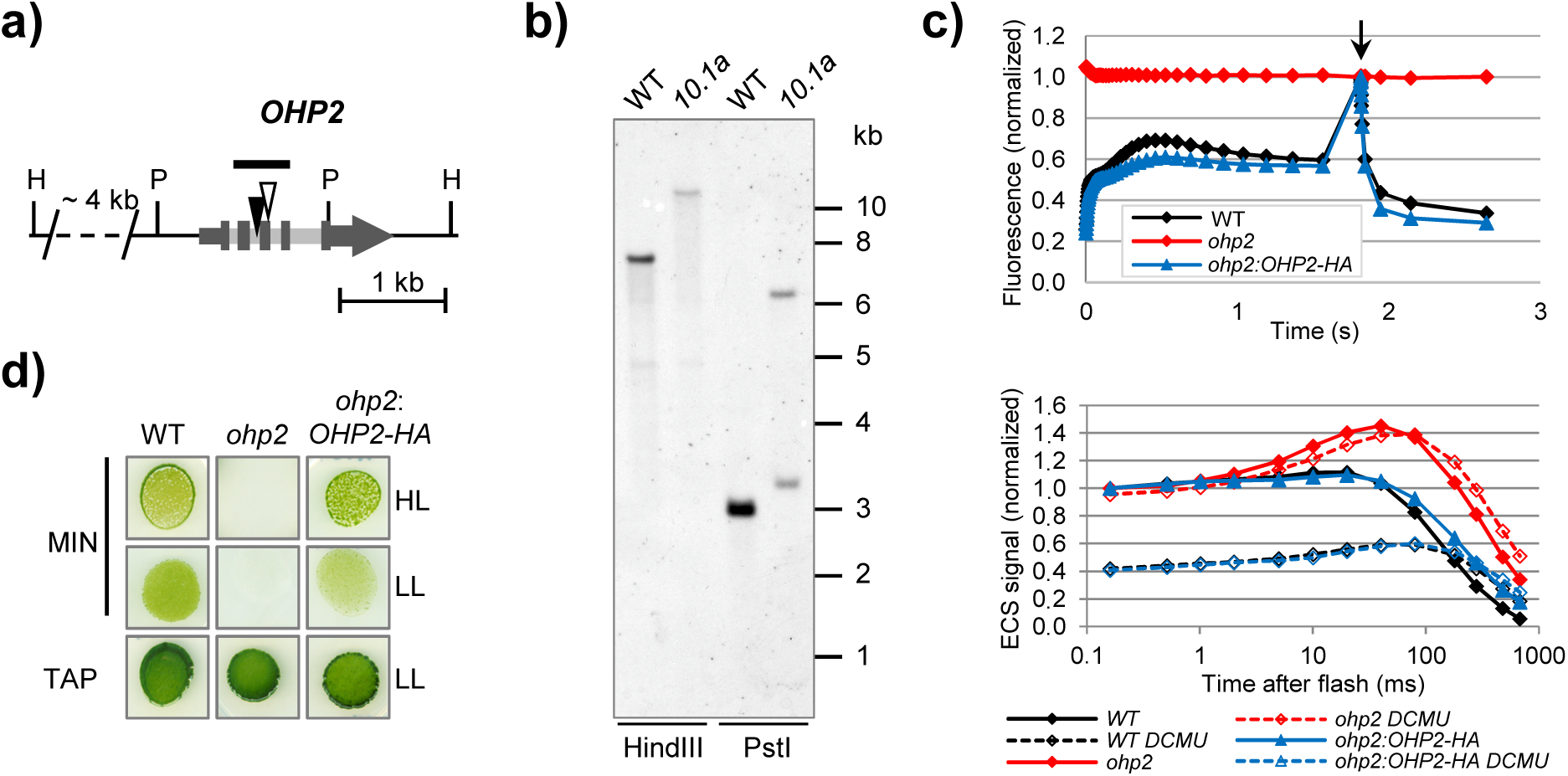
Identification and complementation of the mutation in *OHP2*. a) Schematic representation of the genomic region of *OHP2* (*Cre06.g251150*) on chromosome 6. Grey boxes indicate exons, smaller grey arrow and box 5’ and 3’ UTRs, respectively, and light grey boxes introns. For simplicity, upstream and downstream loci are not displayed. The probe used in b) is shown as black bar above the gene model. The insertion site of a putative >5 kb *TOC1* transposon in exon 3 of the *OHP2* gene in the mutant strain *10.1a* is indicated by an open triangle (compare Supplemental Figure S2). The filled black triangle in intron 2 of the *OHP2* gene indicates the position of a polymorphism caused by an additional putative insertion in the mutant and its background strain as identified during Illumina sequencing (compare Supplemental Figure S2). Given position of HindIII (H) and PstI (P) restriction sites are based on the reference genome of *Chlamydomonas reinhardtii* v5.5 in Phytozome. Note that this sequence is based on the mt+ genome assembly of strain *CC-503* and does not necessarily reflect the restriction site positions in the investigated mt-allele. b) Southern blot analysis of genomic DNA from the Jex4 recipient strain (WT) and the *10.1a* mutant. 10 µg of genomic DNA were fractionated in a 0.8% agarose gel after digestion by restriction enzymes HindIII or PstI, respectively, blotted onto a nylon membrane, and hybridized with the *OHP2*-specific probe indicated in a). c) Photosynthetic parameters of the Jex4 recipient strain (WT), the *ohp2* mutant and *ohp2* complemented with pBC1-CrOHP2-HA (*ohp2:OHP2-HA*, compare construct 2 in Supplemental Figure S3). Fluorescence induction kinetics (upper panel), measured under illumination at 135 µE/m^2^/s, followed by a saturating pulse (arrow) and dark relaxation. Fluorescence intensity is normalized to the Fm value. Electrochromic shift at 520 nm (lower panel), measured in the absence or presence of DCMU and hydroxylamine. Values are normalized to the signal after the saturating flash. d) Growth test. Cells were resuspended in sterile H_2_O to a concentration of 10^5^ cells/mL and spotted onto acetate-containing (TAP) or High Salt Minimum media (MIN) and grown for 6 d under higher light (HL) at 100 µE/m^2^/s or low light (LL) at 30 µE/m^2^/s.

The *OHP2* gene of *Chlamydomonas* encodes the One-Helix Protein 2 (OHP2) belonging to the Chl *a/b* binding protein superfamily (Engelken et al., 2010). OHP2 consists of 144 amino acids with the N-terminal 26 amino acids predicted to represent a chloroplast transit peptide (cTP). The expected mature protein has a molecular mass of 13.4 kDa.

### The *ohp2* mutant reveals a complete loss of PSII activity and a reduced Chl content

As summarized in Supplemental Table S1, fluorescence induction kinetics and electrochromic shift (ECS) analysis of the *ohp2* mutant revealed a complete lack of PSII, while PSI and Cyt*b_6_f* activities appeared normal. PSII quantum yield (Fv/Fm) was negative in the *ohp2* mutant because of the small initial drop in fluorescence, typical of mutants completely lacking PSII (Figure 1C, upper panel). This is due to chemical quenching by plastoquinone brought about by the oxidation of plastoquinol by PSI. Similarly, the PSII/PSI ratio, measured from the amplitude of the initial ECS after a flash in the absence and presence of the PSII inhibitors DCMU and hydroxylamine, indicated loss of PSII charge separation (Figure 1C, lower panel). The small effect of the inhibitor treatment on the *ohp2* mutant was ascribed to hydroxylamine and was also observed in PSII-null mutants such as *psbA-FuD7* (Bennoun et al., 1986). Together, these results revealed that PSII activity is completely abolished in the mutant. When cultivated as liquid culture under photo-heterotrophic conditions, the *ohp2* mutant displayed a lighter-green coloration than the wild type (WT). Determination of the Chl *a* and *b* content showed that, on a per-cell basis, the *ohp2* strain accumulated only about 58% of the total Chl present in the WT. The Chl *a* content decreased more than the Chl *b* level, leading to a decreased Chl *a*/Chl *b* ratio in *ohp2* (Supplemental Table S1), typical of photosynthetic reaction center mutants.

### Complementation of the *ohp2* mutant with *Chlamydomonas* OHP2

Complementation studies were carried out using a plasmid harboring a Pm resistance cassette along with the *OHP2* sequence. We used the full length *OHP2* CDS 3’-terminally fused to a hemagglutinin (HA)-tag encoding sequence, placed under control of the strong nuclear *PSAD* promoter (construct 2 in Supplemental Figure S3). Transformants first selected for Pm resistance were tested for photoautotrophic growth on minimum medium under high or low light. Approximately three quarters of the analyzed clones exhibited restored photoautotrophic growth shown exemplarily for one clone (*ohp2*:*OHP2-HA* in Figure 1D). This strain was chosen for further experiments, with full restoration of photosynthetic capacities, as judged from its WT-like fluorescence induction and ECS kinetics as well as largely restored Chl accumulation (Figure 1C, Supplemental Table S1). These results confirm that the PSII deficient phenotype of the strain is due to inactivation of *OHP2*. They also show that OHP2 can accommodate a small tag at its C-terminus with no deleterious effect on its functionality.

### Functional analysis of the OHP2 protein in *Chlamydomonas*

Mature OHP2 from *Chlamydomonas* contains a single C-terminal transmembrane helix including highly conserved residues required for Chl binding (Figure 2A). Based on topology experiments of Li and colleagues (2019) for the *Arabidopsis* OHP2 protein (AtOHP2), the less conserved N-terminus of OHP2 should face the chloroplast stroma, while the C-terminus is located on the luminal side. In addition, multiple sequence alignment and hydropathy analysis indicate a well-conserved hydrophobic stretch (HS) of nine residues near the C-terminus which is too short to span the lipid bilayer and seems to be missing in cyanobacterial HLIPs like HliD or HliC (compare Figure 2A; Supplemental Figure S4). This HS might be an evolutionary leftover from a second helix of the ancestral two-helix Stress-Enhanced Proteins (SEP) from which OHP2 potentially originates (Andersson et al., 2003; Beck et al., 2017; Heddad et al., 2012).

**Figure 2.**
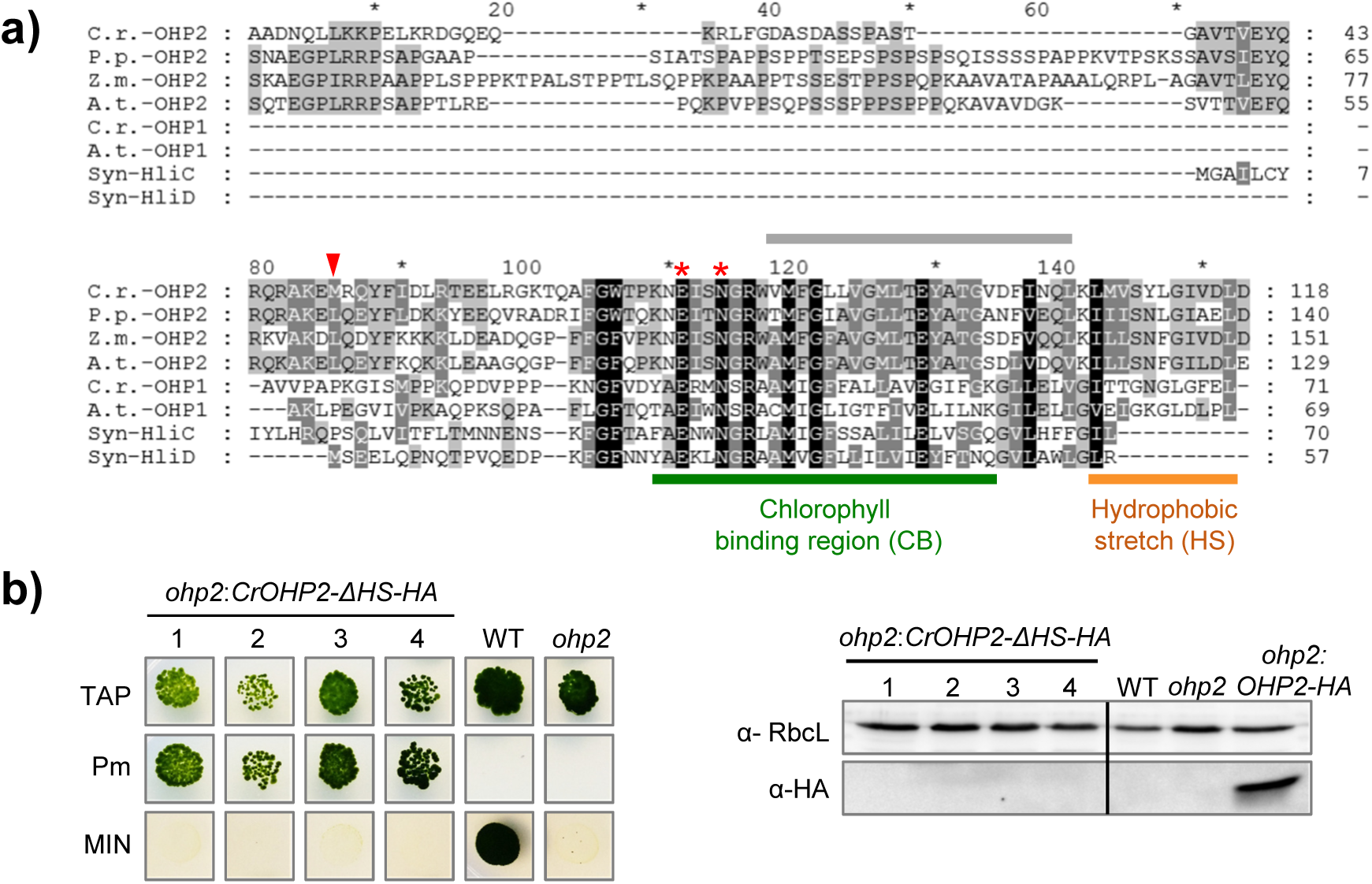
Protein sequence alignment and complementation analysis. a) Sequence alignment of eukaryotic One-Helix Proteins OHP1 and OHP2 and two cyanobacterial HLIPs. OHP2 and OHP1 protein sequences from C.r. (*C. reinhardtii*, v5.5, *Cre06.g251150* and *Cre02.g109950*), A.t. (*Arabidopsis thaliana*, TAIR10, *AT1G34000.1* and *AT5G02120*), P.p. (*Physcomitrella patens*, v3.3, Pp3c2_26700V3.1), and Z.m. (*Zea mays*, PH207 v1.1, Zm00008a032025_T01) were obtained from Phytozome (https://phytozome.jgi.doe.gov/pz/portal.html). Amino acid sequences of the High light-induced proteins HliC (*ssl1633*) and HliD (*ssr1789*) from *Synechocystis sp.* PCC6803 were taken from CyanoBase (http://www.kazusa.or.jp/cyano/). The multiple sequence alignment was performed by using ClustalW (Thompson et al., 2002), manually edited and displayed with Genedoc (Nicholas et al., 1997). Black shading represents 100% conservation, dark grey and grey 60% and 40%, respectively. The positions of the predicted Chl binding region (CB) as well as a hydrophobic stretch (HS) at the C-terminus are indicated (compare Supplemental Figure S4A). Two residues described to be important for Chl binding in LHCB from spinach are labeled with red asterisks (Kühlbrandt et al., 1994). The N-terminal chloroplast transit peptides predicted by TargetP-2.0 (Emanuelsson et al., 2000; Nielsen et al., 1997) for all eukaryotic proteins shown, are not included in the alignment. The *Chlamydomonas* OHP2-based prediction of a transmembrane helix by TMPred (Hofmann & Stoffel, 1993; https://embnet.vital-it.ch/software/TMPRED_form.html), is indicated as a grey bar above the sequence (compare Supplemental Figure S4B). The mutation identified in the *Chlamydomonas OHP2* gene corresponds to residue M76 of the protein (indicated by a red triangle) which lies within the fully-conserved stretch in the N-terminal part of the protein. b) The C-terminal hydrophobic stretch is required for restoration of photoautotrophy. Growth test (left panel). The *ohp2* mutant was transformed with the construct pBC1-CrOHP2-ΔHS-HA (see construct 5 in Supplemental Figure S3). Pm-resistant transformants (*ohp2*:*CrOHP2-ΔHS-HA*) were tested for photoautotrophic growth on HSM plates (MIN). Immunoblot analysis (right panel) of Pm-resistant transformants. 40 µg of total proteins were separated on 15% denaturing polyacrylamide gels and analyzed with the antibodies indicated on the left. The complemented strain *ohp2*:*OHP2-HA* was used as positive control for the accumulation of HA-tagged proteins. Samples were run on the same gel but not in adjacent lanes as indicated by a vertical black line.

To test the functional role of this C-terminal hydrophobic region we transformed the mutant with a construct for the expression of a truncated OHP2 protein (pBC1-CrOHP2-ΔHS-HA, construct 5 in Supplemental Figure S3) where the last 14 amino acids are replaced by the HA-tag. Thirty transformants selected on Pm were tested for photoautotrophic growth on minimal medium, but none showed restoration of photoautotrophy (Figure 2B, left panel). No accumulation of the truncated OHP2-HA could be detected in the transformants by immunoblots using an α-HA-antibody (Figure 2B, right panel). These results suggest that even though cyanobacterial orthologs do not have the HS, the last 14 amino acids are essential for OHP2 stability in *Chlamydomonas*.

A construct for the expression of a fusion of full-length OHP2 with GFP, adapted to the codon usage of *Chlamydomonas* (CrGFP; construct 3 in Supplemental Figure S3), yielded Pm-resistant transformants, but none was photoautotrophic. Presumably, the large soluble GFP-tag interferes with the function or proper localization of OHP2. However, when we directly fused the predicted transit peptide of OHP2 to CrGFP (construct 4 in Supplemental Figure S3), we observed GFP protein accumulation in two out of eight immunologically analyzed clones upon transformation of the *UVM4* strain which is known for its high capacity to express nuclear transgenes (Supplemental Figure S5A, upper panel; Neupert et al., 2009). Confocal laser scanning microscopy of one transformant showed that the GFP fluorescence signal co-localized with the Chl autofluorescence in the single cup-shaped chloroplast (Supplemental Figure S5A, lower panel), clearly distinct from the cytosolic control (CrGFP). Moreover, immunological analysis of cellular subfractions of the strain *ohp2:OHP2-HA* described above revealed a membrane localization of the ~14 kDa OHP2-HA tagged protein comparable to that of the integral thylakoid membrane protein D2 (Supplemental Figure S5B).

Taken together, these results indicate that OHP2 is a thylakoid membrane-localized protein. Moreover, while a C-terminal fusion of a small tag to OHP2 still allows the protein to fulfil its native function, removal of its C-terminus or introduction of a large GFP tag severely impairs its function and /or stability.

### Reduced PSII subunit accumulation and synthesis in the *ohp2* mutant

As judged from the analysis of photosynthetic parameters in the *ohp2* mutant, in particular the lack of effect on PSI, OHP2 appears exclusively involved in PSII biogenesis. Accordingly, immunoblot analyses revealed no change in the accumulation of the large subunit of Rubisco (RbcL), the PSI reaction center protein PsaA or Cyt*f* of the Cyt*b_6_f* complex (Figure 3A). In contrast, the PSII core proteins D1 and D2 were below the detection limit. Typical for mutants lacking D1 or D2, the core antenna protein CP43 accumulated to some extent, about 12% of the level in WT (Figure 3A). Complementation of *ohp2* with the *OHP2* cDNA fully restored protein accumulation in the strain *ohp2:OHP2-HA* (Figure 3A).

**Figure 3.**
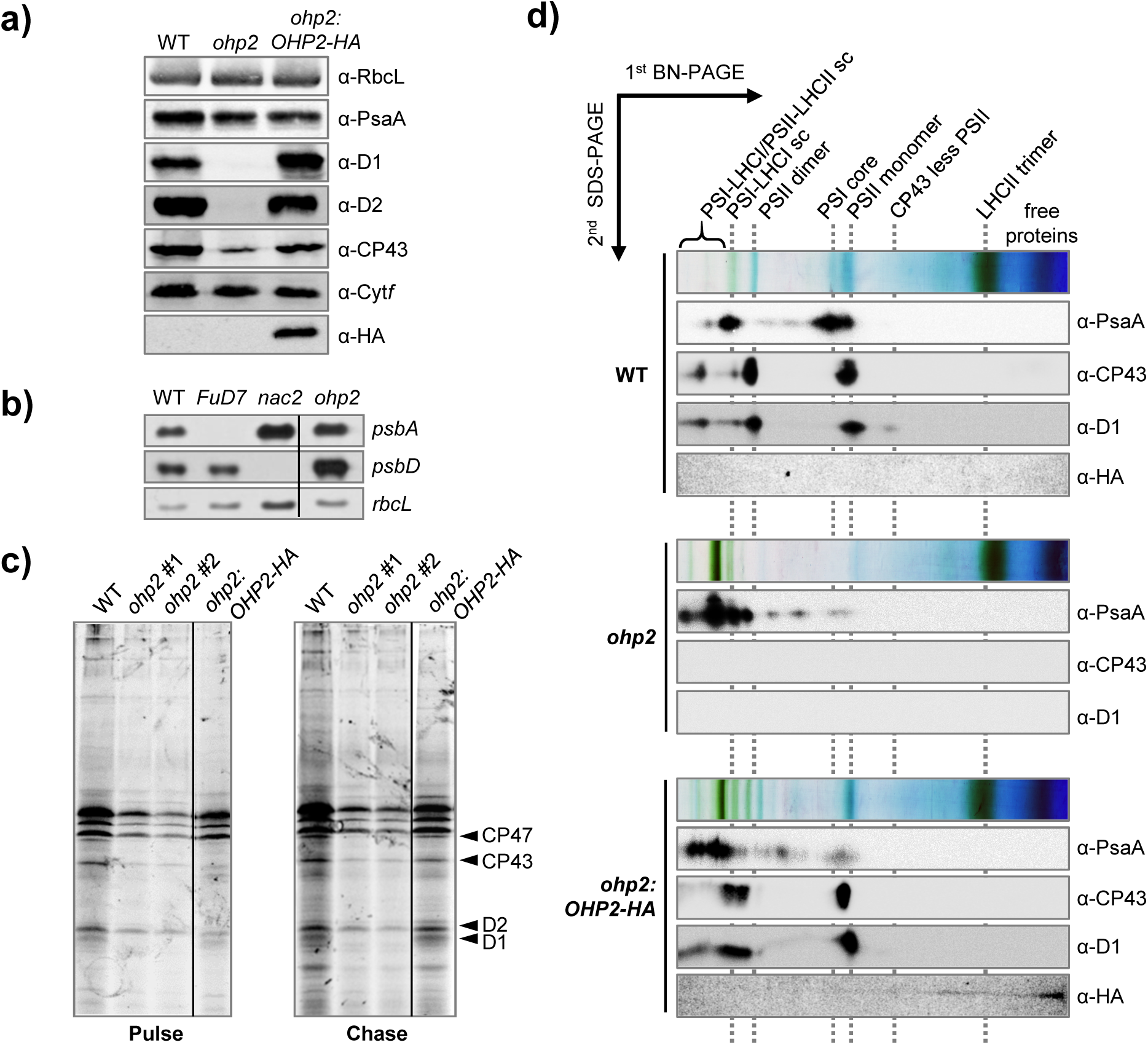
Drastically diminished D1 protein accumulation and synthesis in the *ohp2* mutant. The following strains were subjected to analysis: Jex4 (WT), *ohp2* and the complemented strain *ohp2:OHP2-HA* clone *#53 (ohp2:OHP2-HA)*. a) Accumulation of photosynthesis related chloroplast proteins. 30 µg of total proteins from indicated strains were separated by 12% (α-D1, α-PsaA, α-Cyt*f,* α-CP43, α-D2) or 15% (α-HA, α-RbcL) denaturing polyacrylamide gels and analyzed by the antibodies indicated. b) Accumulation of photosynthesis related chloroplast transcripts. 3 µg of total cellular RNA from indicated strains were fractionated by denaturing agarose gel electrophoresis and blotted onto a nylon membrane. Membranes were hybridized with probes specific for *psbA* and *psbD*. For loading control, the same blot was hybridized with a probe specific for *rbcL* transcripts. *nac2* and *FuD7* mutants were employed as negative controls for *psbD* or *psbA* mRNA accumulation, respectively. *ohp2* was run on the same gel but not in adjacent lanes as indicated by a vertical black line. c) Synthesis of PSII subunits. ^14^C labelling of chloroplast-encoded proteins was performed as described previously in Spaniol et al. (2021). Autoradiogram of cells pulsed for 5 min with ^14^C-acetate in the presence of cycloheximide (left), then chased for 45 min after washing and chloramphenicol addition. For *ohp2,* two independently grown cultures (#1, #2) were analyzed. The complemented strain was run on the same gel and exposed in the same conditions, but in non-adjacent lanes as indicated by a vertical black line. d) 2D BN-PAGE analysis of photosynthetic protein complexes in indicated strains. After solubilization of membranes with 1.5% (w/v) dodecyl-b-D-maltopyranoside, thylakoid proteins were separated by 5 to 12 % BN gels in the first dimension and 12% SDS-gels in the second dimension. Photosynthetic complexes were detected using the indicated antibodies. The positions of major PSI and PSII complexes are designated. sc: supercomplex

Northern blot analysis showed no significant alteration of *psbA* and *psbD* transcript accumulation in *ohp2* compared to the WT (Figure 3B). To determine whether the mutation instead affects translation or prevents stabilization of chloroplast-encoded PSII subunits, ^14^C pulse chase labeling experiments were carried out in the presence of cycloheximide (Figure 3C). Two independently grown *ohp2* cultures revealed a reduction of radiolabeled CP47, and to a lower extent of CP43 and D2 proteins in comparison to the WT. However, the most significant effect was observed for the D1 protein for which no incorporation of the radiolabel was detected, neither during the pulse nor after the chase. This suggests that the primary defect in the *ohp2* mutant is either a deficiency in D1 synthesis or a very fast degradation of newly synthesized D1. The effects on other PSII subunits may be ascribed to assembly-dependent translational control (CES; Choquet and Wollman, 2009) or reduced stability of the unassembled subunits.

Our efforts to raise a functional antibody against recombinant OHP2 having failed, we used the *ohp2:OHP2-HA* strain, along with the WT and *ohp2*, to examine the assembly of PSII and the location of OHP2. Crude membrane fractions were solubilized and subjected to 2D-BN/SDS-PAGE (Figure 3D). As anticipated from the unaffected accumulation of the PSI subunit PsaA (Figure 3A), all PSI-related complexes, including PSI core and supercomplexes, were assembled in the *ohp2* mutant. Remarkably, a higher ratio of PSI supercomplexes to PSI core complexes was observed in the *ohp2* mutant, suggesting that LHCI antenna in this strain is more tightly coupled to the PSI core than in the WT. This phenomenon was only partially reverted in the complemented strain. All PSII complexes formed in the WT were below the detection limit in the absence of the OHP2 protein. In agreement with the restored photoautotrophic growth and PSII protein accumulation in the complemented strains, all PSII related complexes were present in the analyzed *ohp2:OHP2-HA* strain. Interestingly, different from *Arabidopsis,* where AtOHP2 was detected as a distinct signal at the size of a ~150 kDa PSII RC-like complex (Li et al., 2019), the HA-tagged OHP2 protein in our complemented strain was mainly detected in the low molecular range, reaching up as a weak smear to the first monomeric CP43-less PSII RC47 subcomplex (Figure 3D). This suggests that in *Chlamydomonas* the putative OHC does not withstand the solubilization conditions used for BN-PAGE. Accordingly, we were unable to pull-down OHP2 interactants using the HA-epitope as a bait.

### Ribosome profiling reveals that D1 is still translated in the absence of OHP2

Based on the molecular characterization of the *ohp2* mutant, the major effect of the loss of OHP2 in *Chlamydomonas* seems to be the missing accumulation of the PSII reaction center protein D1, in line with observations in other organisms. However, our ^14^C pulse labelling experiments (Figure 3C) can be interpreted either as an absence of translation of the *psbA* mRNA or as a very rapid or even co-translational degradation of newly synthesized D1. To address this question and obtain a quantitative image of chloroplast translation in the *ohp2* mutant, we employed a previously established targeted chloroplast ribosome profiling approach (Trösch et al., 2018). Ribosome footprints (FP) from WT and mutant were extracted and analyzed by hybridization of highly tiled microarrays covering all open reading frames of the chloroplast genome. In parallel, total RNA was isolated, fragmented and detected by the same approach. Direct correlation between the three biological replicates showed a high reproducibility both for determining RNA accumulation (r > 0.96) and translation output (r > 0.93) (Supplemental Figure S6A, Supplemental Dataset 1).

General gene expression defects were examined by averaging all probe intensities for mRNA or ribosome FPs that covered individual chloroplast CDS, respectively. Direct plotting of mRNA intensities showed highly comparable RNA abundances between the WT and the mutant (r = 0.98, Figure 4A). This suggests that the lack of OHP2 is not causing any obvious transcription or RNA stabilization defects. In contrast, the translational output, as determined by relative FP abundance for each ORF, revealed clear differences between mutant and WT cells and hence, reduced correlation (r = 0.9, Figure 4A).

**Figure 4.**
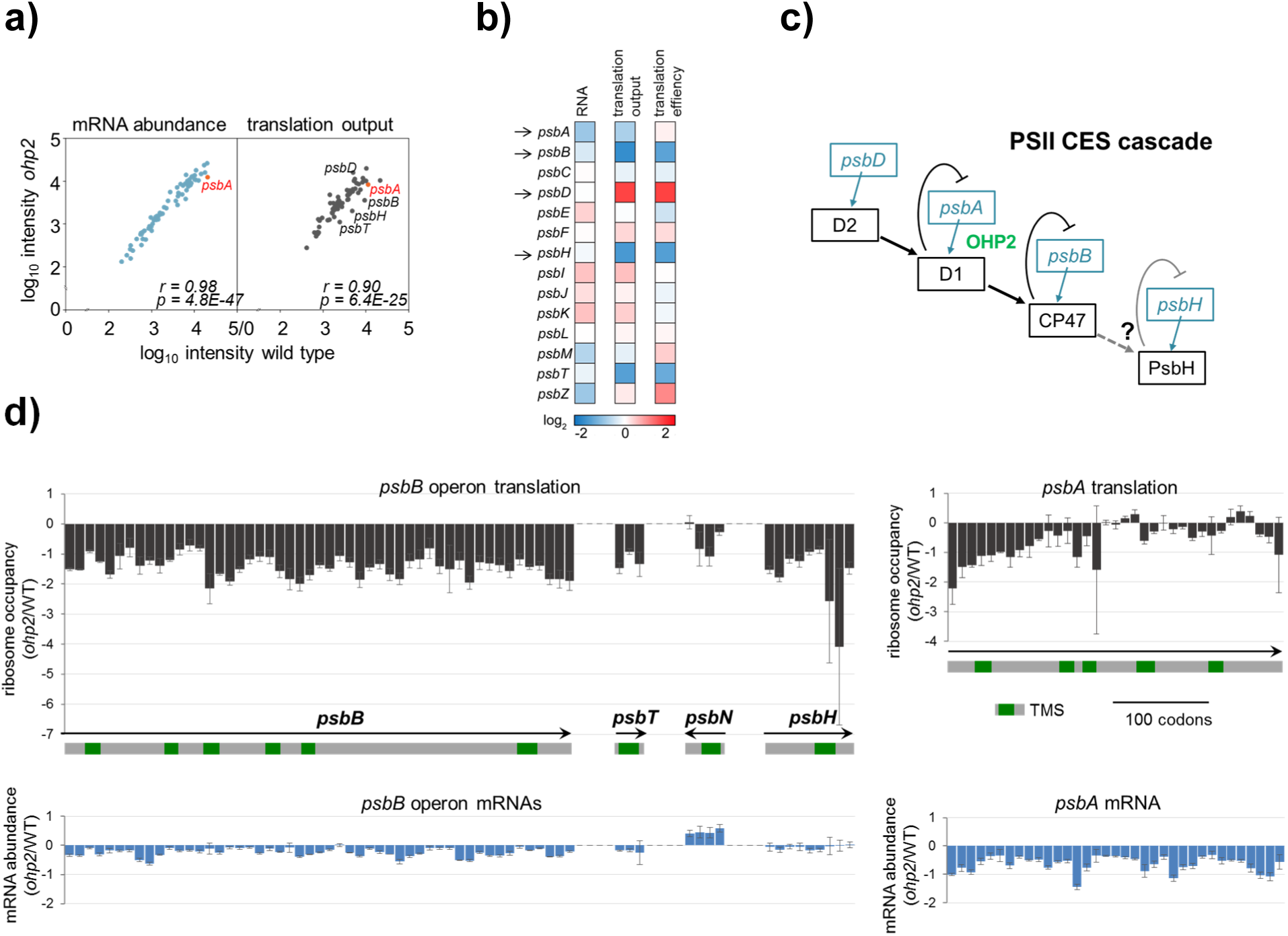
Targeted ribosome profiling of chloroplast translation reveals protein synthesis defects in the *ohp2* mutant. Ribosome profiling and transcript analysis of WT and *ohp2* mutant grown mixotrophically under 30 µE/m^2^/s. a) The average mRNA (blue) and ribosome FP (dark grey) abundances were calculated from three independent biological replicates and plotted in log_10_ scale for *ohp2* mutant versus WT, respectively. mRNAs encoding PSII subunits, which display altered translation in the mutant are highlighted. *Pearsons’s r*-value and *p*-value are given in *n*E*m* non-superscript format for *n•*10*m*. b) The relative average transcript abundances (RNA), translation output and translation efficiency were calculated for each chloroplast reading frame in both *Chlamydomonas* strains, normalized to overall signal intensities, and plotted as heat map (*ohp2* versus WT) in log_2_ scale. Increased RNA accumulation, translation output or translation efficiency in the mutant is shown in red, reduced levels are shown in blue (see scale bar). PSII subunits that are further discussed are highlighted with an arrow. c) CES cascade of Photosystem II. When newly synthesized CES polypeptides, like D1 and CP47, cannot assemble, they repress the translation initiation of their encoding mRNA as described before by Minai and coworkers (2006). Reduced translation of *psbH* may indicate that it represents a further component of the CES cascade. d) For the Open Reading Frames (ORFs) of the *psbB* operon and *psbA* (introns not shown), normalized ribosome FP intensities were plotted as mean log_2_ ratios between *ohp2* mutant and WT. Error bars denote differences between three independent biological replicates. Grey bars below the panels indicate respective ORFs with trans-membrane segments (TMS in green). Related to Supplemental Figures S7-S9 and Supplemental Dataset 1.

Unexpectedly, averaged FP intensities of *psbA* showed only a mild reduction in the mutant (Supplemental Dataset 1). FP abundance was reduced by 1.6-fold (+/− 0.16), which combined with a slight reduction in mRNA abundance, led to a practically unchanged translation efficiency (Figure 4B). For a more detailed view, we also plotted ribosome occupancy (log_2_ of the *ohp2*/WT ratio) over the *psbA* ORF (Figure 4C, left panel). A more pronounced reduction (up to 4-fold) in ribosome occupancy in *ohp2* can be observed over the first half of the ORF (~160 codons, up to the 3^rd^ transmembrane segment of nascent D1). In the second half, only mild reduction or even a slight increase in ribosome occupancy could be detected. For other PSII subunits, the most pronounced effect was the reduced translation of CP47 (PsbB), PsbH, and PsbT, with up to 3-fold lower translation output for *psbB* (Figure 4B and Supplemental Figure S6B). This observation agrees with the hierarchical CES cascade contributing to the biogenesis of PSII in *Chlamydomonas*, where the presence of D2 is required for high-level translation of D1, which in turn is a prerequisite for efficient translation of the core antenna protein CP47 (Figure 4D; reviewed in Choquet and Wollman, 2009). When assembly of D1 and CP47 subunits is compromised, the unassembled proteins repress the translation initiation of their encoding mRNA (Minai et al., 2006). Also, in accordance with previous investigations of PSII mutants (de Vitry et al., 1989), we observed no effect on translation of the second inner antenna protein CP43, encoded by the *psbC* mRNA, confirming that its rate of synthesis is not dependent on the assembly with other PSII subunits.

Interestingly, also PsbH, which only recently has been hypothesized to be part of the CES cascade downstream of D1 (Trösch et al., 2018), exhibited a clearly reduced translation output in *ohp2,* thus further supporting its role as a CES subunit (Figures 4A, 4B, 4D). A comparable effect was seen for the translation of *psbT.* Remarkably, the three affected genes are encoded in the *psbB-T-H* operon. To obtain a more detailed view on the translation of this polycistronic transcript, we further looked at relative ribosome FP accumulation along this *psbB-T-H* operon, calculated as mean log_2_ ratios to directly compare mRNA abundance and ribosome occupancy between mutant and WT. While the RNA of the operon was only marginally reduced, a clear and relatively even reduction of ribosome occupancy was seen for all ORFs (Figure 4C, right panel). It is noteworthy that also *psbN*, which is positioned between *psbT* and *psbH* in antisense orientation, exhibits a reduced translational efficiency while the mRNA levels appeared slightly increased. PsbN acts as an assembly factor for PSII but is not part of the final complex (Knoppová et al., 2022).

Complementary dynamics were observed for the synthesis of D2, which showed increased ribosome FP intensities over the entire ORF in *ohp2* while maintaining constant mRNA abundance (Figures 4A, 4B, Supplemental Figure S7). As D2 is the dominant CES subunit (Figure 4D), this likely points to a compensatory effect caused by the lack of functional PSII. Similarly, other translational alterations in *ohp2*, like the reduction of ribosome FPs for the large Rubisco subunit RbcL or an upregulation of all chloroplast-encoded ATP synthase subunits (Supplemental Figure S6), may rather be secondary effects caused by a diminished photosynthetic electron flow and/or energy limitation.

### Proteomic analysis reveals the presence of HCF244 in the *ohp2* mutant

For a quantitative and comprehensive view on the proteome composition in the *ohp2* mutant, proteomic shot-gun analysis was conducted on whole cell fractions (Figure 5A, Supplemental Dataset 2). The analysis additionally included an *ohp2* strain in which the PSII phenotype was genetically suppressed. Such photoautotrophic suppressor strains occurred at a high frequency (approximate rate: 5.5 × 10^−6^) under conditions selecting for photoautotrophic growth. Illumina sequencing and Southern blot analysis of photoautotrophic *ohp2* derivatives showed that they still contained the *TOC1* transposon in *OHP2* (Supplemental Figures S2, S8A), indicating suppression of the PSII phenotype by a second site mutation. The suppressor strains showed partially restored photosynthetic parameters (Fv/Fm ratios of 0.46-0.64), a re-accumulation of PSII subunits indicated by restored D2 levels, and the ability to grow photoautotrophically in liquid medium under strong illumination (Supplemental Figures S8B-D). We proteomically analyzed suppressor strain *M-Su1* to distinguish the direct effects of the lack of OHP2 from those linked to PSII loss. In addition, LC-MS analysis was carried out on membrane fractions of WT, *ohp2* and the suppressed strain (Figure 5B, Supplemental Dataset 3). These two analyses allowed reliable quantification of 3065 and 1127 proteins, respectively, with very high reproducibility between the biological replicates (Supplemental Figure S9A). The complemented strain *ohp2:OHP2-HA* was included as a control, and showed that most of the alterations observed in mutant cells were reverted by complementation (Supplemental Figure S9B).The most obvious defect observed in the *ohp2* strain, apart from the expected absence of OHP2 itself, was a severe depletion of the majority of PSII proteins. The most dramatic effect was on D1 showing a ~200/640-fold reduction in whole cell and membrane fractions, respectively. However, the detection in the *ohp2* mutant of five D1 peptides arising from the C-terminal third of the protein supports the notion, based on the results of the ribosome footprinting, that the *psbA* CDS is translated over its entire length. For the other large PSII subunits D2 and CP47, the depletion was less pronounced. The least affected subunit was CP43, in accordance with results obtained with other PSII mutants (de Vitry et al., 1989). The small molecular weight subunits of PSII were difficult to assess in these experiments, but we noticed a massive depletion of PsbH in the mutant (Supplemental Datasets 2, 3), in agreement with the ribosome profiling results. Other PSII subunits were severely depleted, at least at the membrane level, including the newly-discovered algae-specific subunit PBAS1 (= PBA1, Putatively Photosystem B Associated 1; Spaniol et al., 2021). Among the extrinsic luminal Oxygen-Evolution Enhancer (OEE) subunits, no significant change was observed in whole cells, but OEE1 was depleted in membranes, in line with previous reports (de Vitry et al., 1989).

**Figure 5.**
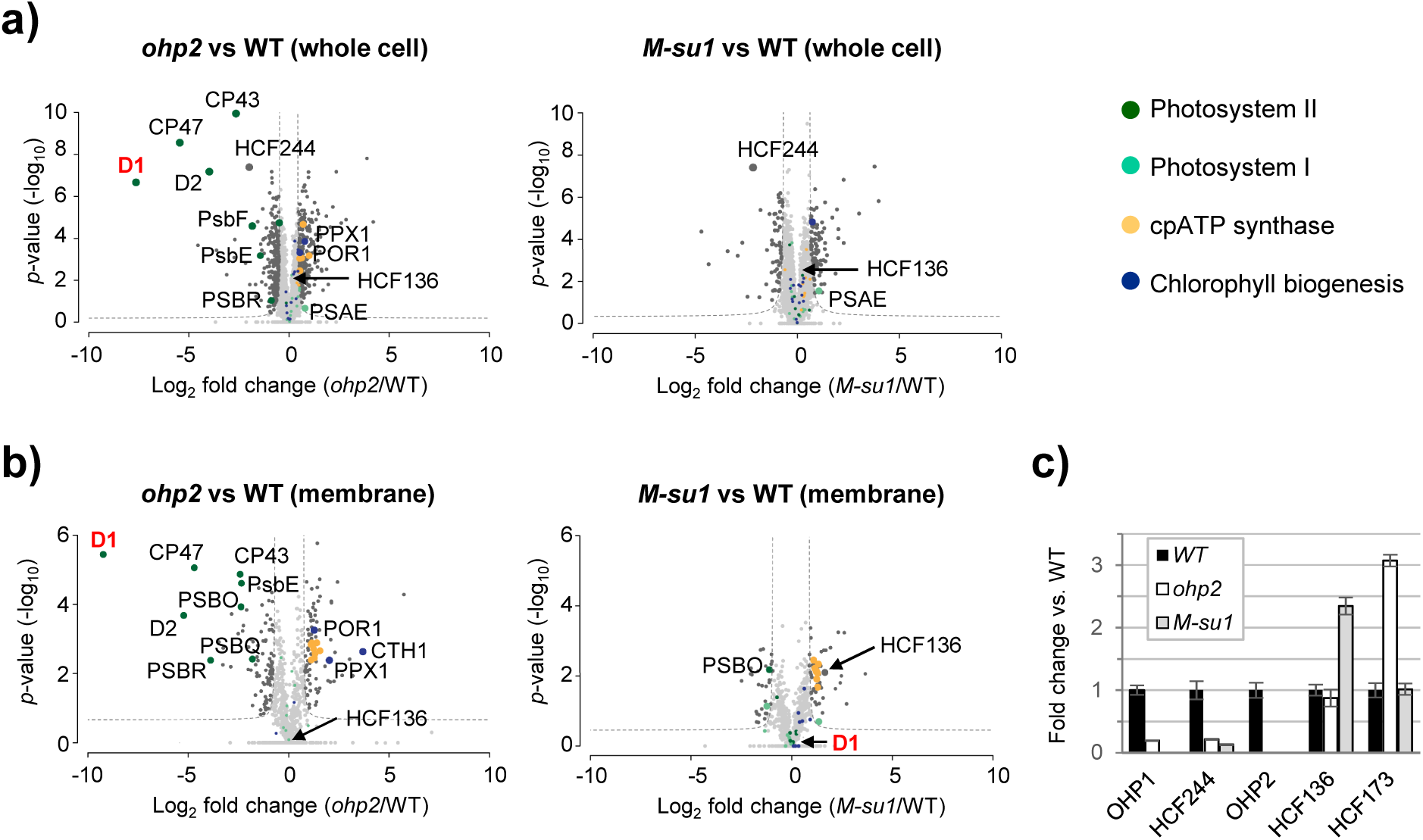
Altered proteome composition in *ohp2* mutants and suppressor strains. Volcano plots representing the relative proteome changes of whole cell lysates (a) of *ohp2* mutants versus WT (left), suppressor line 1 versus WT (right) and thylakoid membrane proteins (b) as comparison between *ohp2* mutant versus WT (left) and suppressor line 1 versus WT (right). For b), proteins of the membrane fraction were enriched by crude fractionation. All experiments were performed in at least three independent biological replicates. Mean fold change of LFQ values (in log_2_) is plotted on the x-axis, *p*-values (in −log_10_) are plotted on the y-axis. Light grey dots represent proteins with no significant change, dark grey dots show proteins that are significantly different with FDR <0.05 and S_0_=1. Proteins of PSI, PSII, the chloroplast ATP synthase, and of proteins involved in Chl biogenesis are marked in color as depicted in the legend on the right. Large colored dots are significantly different. c) Abundance of selected proteins in membrane protein extracts as determined by targeted mass spectrometry in different strains as compared to WT. Error bars indicate standard deviation over three biological replicates. Missing values indicate instances where no peptides could be found or are present below baseline signal of app. 5% compared to WT as judging from dotp values and fragment coelution profiles.

Of particular interest for us was the presence in the mutant and suppressor strains of OHP1 and HCF244, the partners of OHP2 in the plant OHC. Probably due to its small size and hydrophobicity, OHP1 was detected neither in the whole cell extracts nor in the untargeted analysis of membrane fractions, even in the WT. To further refine our analysis, the same membrane fractions were also subjected to a targeted proteomics approach, whereby specific peptides were monitored at high resolution. By reducing interferences from the MS1 level, this allows a more reliable quantitation. Five proteins were selected: OHP2, OHP1, HCF244, HCF136 and HCF173, using a total of 25 peptides (Figure 5C and Supplemental Dataset 4). This targeted analysis used two peptides for OHP1 and showed a low signal in membrane fractions of the *ohp2* mutant (18% of WT). This suggests that in *Chlamydomonas*, OHP1 is not completely dependent on OHP2 for its stabilization. The OHP1 signal was not detected in the suppressed strain.

In whole cell extracts, HCF244 accumulated to ~25% of the WT level in *ohp2* and in the suppressor strain (Supplemental Dataset 2). Like OHP2 accumulation, this was largely complemented by expression of the *OHP2-HA* transgene. Untargeted analysis of the membrane fractions revealed the presence of HCF244 in the WT but not in *ohp2* or the suppressed strain. However, the more sensitive targeted analysis, based on eight peptides, revealed the presence of HCF244 in the membrane fractions of the mutant and suppressor strains, with an abundance of 22% and 13% relative to WT levels, respectively (Supplemental Dataset 4, Figure 5C). Together, these results indicate, that in contrast to *Arabidopsis*, the presence of OHP2 is not absolutely necessary for HCF244 accumulation, nor for its association with the membrane.

### Proteomic analysis of other PS subunits, assembly factors and chloroplast enzymes

We also examined the accumulation of other known PSII biogenesis factors, in search of secondary effects of the lack of OHP2, or possible mechanisms or consequences of suppression (Figure 5; Supplemental Datasets 2 - 4). At the whole cell level, the abundance of HCF173 was not affected in the mutant or suppressor, but targeted analysis of the membrane fractions revealed a ~3-fold higher level in *ohp2*, compared to the WT or suppressed strains, suggesting a regulatory mechanism. Another known interactant of the *psbA* mRNA in plants is SRRP1: its abundance was significantly increased in the *ohp2* mutant cells and restored to normal by complementation. In the suppressor strain, SRRP1 was barely detectable. Similarly, the *psbA* translation activator TBA1 (Somanchi et al., 2005) overaccumulated in the mutant.

The luminal protein HCF136 is an early interactant of the D2/Cyt*b*_559_ pre-complex, necessary for assembly of pD1 into the RC (Plücken et al., 2002). Its abundance in whole cells was not significantly altered, but both the untargeted and targeted analyses of the membrane fractions revealed a significantly higher membrane association in the suppressor strain (~3-fold when compared to the WT; Figure 5; Supplemental Datasets 2 - 4). This may reveal a more prolonged association of HCF136 with the RC when the absence of OHP2 slows down the biogenesis process.

PSB28 binds to CP47 on the stromal surface and distorts the Q_B_-binding pocket, as revealed by the structural analysis of a cyanobacterial PSII assembly intermediate also comprising PSB27, which binds to CP43 and the transmembrane protein PSB34 on the luminal side (Xiao et al., 2021; Zabret et al., 2021). Here, PSB28 and PSB27 showed interesting contrasted behaviors. At the whole cell level, the accumulation of PSB28 showed no significant change between strains. However, it was undetectable in the membrane of the *ohp2* strain or of the suppressor. In contrast, PSB27 was overaccumulated in *ohp2* cells, with no change in its abundance in membrane fractions. PSB27 was also somewhat overaccumulated in the suppressor, this time accompanied by a marked increase in its association with the membrane. We also noticed an increased accumulation in the *ohp2* mutant of PSB33/TEF5, a factor involved in the interaction of photosystems and antenna complexes (Fristedt et al., 2015; Nilsson et al., 2020), suggesting stabilization byt prolonged interaction with the antenna in the absence of RC. The mutant also overaccumulated APE1, initially described as involved in the adaptation to high light (Walters et al., 2003) and later found as a specific interactant of *Arabidopsis* OHP1 in pull-down assays (Myouga et al., 2018) and associated with the RCIIa complex isolated from a Synechocystis strain lacking YCF39/HCF244 (Knoppová et al., 2022).) Overaccumulation was also observed for the homolog of Slr1470, another component of the RCIIa complex (Knoppová et al., 2022). This suggests that absence of the OHC in *Chlamydomonas* leads to prolonged association (and stabilization) of those assembly factors whose interaction with the RC is antagonized by that of the OHC. Finally, we observed increased membrane association of PSB29/THF1 in the mutant, which was partially reverted in the suppressor strain. THF1 has been shown to be involved in the dynamics of PSII-LHCII supramolecular organization and to associate with the FtsH protease (Becčková et al., 2017; Huang et al., 2013; Wang et al., 2004).

In contrast to PSII, no significant effect was observed on the accumulation of PSI RC subunits: the slight overaccumulation of PSAE in whole cells of the mutant was also found in the suppressed and complemented strains and may therefore be unlinked to OHP2. More interestingly, however, all the LHCI subunits showed increased accumulation in the *ohp2* mutant as well as in the suppressor strain, suggesting a direct effect of the lack of OHP2 (Supplemental Dataset 2). While subunits of the Cyt*b_6_f* complex appeared unaffected in the *ohp2* mutant, we noticed an overaccumulation of those of the ATP synthase (Figure 5A, left panel). This agrees with the increased translational output for the ATP synthase ORFs seen in our ribosome profiling analysis (Supplemental Figure S6B). Overaccumulation of the ATP synthase was fully reversed in the complemented strain (Supplemental Figure S9B). We note, however, that the suppressor strain, although photoautotrophic, still showed some overaccumulation of the ATP synthase in its membrane fraction (Figure 5B). The Rubisco subunits RbcL and RBCS2, in contrast, appeared to accumulate at lower levels in the mutant in whole cells fractions (Supplemental Dataset 2). The significance of this observation is unclear, as their accumulation was not restored to normal in the suppressor strain.

Several enzymes of the porphyrin biosynthesis pathway showed changes in their accumulation or association with the membrane (Supplemental Datasets 2, 3, Figures 5A, 5B). The Chl synthase CHLG, whose cyanobacterial homolog interacts with HliD/Ycf39 (Chidgey et al., 2014), remained unaltered in the *Chlamydomonas* mutant, but other Chl synthesis enzymes showed increased abundance in the membrane fractions. This includes the two subunits of the Mg-chelatase (ChlH, ChlI), the Mg protoporphyrin IX S-adenosyl methionine O-methyl transferase ChlM, the cyclases CRD1 and CTH1 and the protochlorophyllide *a* oxidoreductase POR1. Except for POR1, this increase was not observed at the whole cell level, suggesting that the cell responds to the absence of the OHC complex by stabilizing the interaction between these Chl pathway enzymes and the thylakoid membrane. This effect was largely or completely reversed in the suppressor, pointing to a regulatory mechanism compensating for the impaired stabilization of D1. In the upstream part of the pathway, shared with heme biosynthesis, several enzymes overaccumulated in whole cells of the mutant (Figure 5A): the delta-aminolaevulinic acid dehydratase (ALAD1), the uroporphyrinogen III synthase (UPS1), one of the three uroporphyrinogen III decarboxylases (UPD1), and the protoporphyrinogen oxidase (PPX1). PPX1 abundance also increased in the membrane fraction. The protein FLUORESCENT (FLU), which is involved in the regulation of the whole pathway (Falciatore et al., 2005) and which was severely reduced in a virus-induced *OHP2* gene silenced *Arabidopsis* line (Hey and Grimm, 2018), was not significantly affected in the *Chlamydomonas* mutants.

### The *Arabidopsis* OHP2 complements the *Chlamydomonas ohp2* mutation

The observed phenotypical differences between *ohp2* mutants from higher plants and *Chlamydomonas*, as well as the divergence of the amino acid sequences (Li et al., 2019; Myouga et al., 2018) led us to test whether the *Arabidopsis* protein could complement the *Chlamydomonas* mutant. The N-terminal region of the mature OHP2 proteins is much less conserved than the transmembrane and C-terminal regions and similarity between the *Chlamydomonas* and *Arabidopsis* proteins starts only at residue 65 of the *Chlamydomonas* OHP2 preprotein with an overall identity of 58% (compare Figure 2A).

We used a construct (pBC1-TP_CrOHP2_-AtOHP2-HA, construct 6 in Supplemental Figure S3) fusing the predicted cTP of OHP2 from *Chlamydomonas* with a codon-adapted sequence encoding the mature part of AtOHP2 (Supplemental Data S2). After transformation of the *Chlamydomonas ohp2* mutant and selection for Pm-resistance, 17 of the 20 clones (85%) showed restoration of photoautotrophy and Fv/Fm values of 0.62 – 0.74 (Supplemental Figure S10A). For all of the immunologically analyzed clones, a protein with the expected size of ~ 15 kDa was detected using the α-HA-tag antibody and D1 proteins accumulated to WT levels (Supplemental Figure S10B). Taken together these data indicate WT level PSII accumulation and successful complementation of the *Chlamydomonas* mutant with the *Arabidopsis* protein.

### Slowing down D1 degradation in the *ohp2* mutant can partially restore light-sensitive PSII activity, but not photoautotrophy

The repair of photodamaged PSII involves the selective proteolytic degradation and replacement of the damaged D1 polypeptide with a newly synthesized one (reviewed in Kato and Sakamoto, 2009; Nixon et al., 2010). The thylakoid protease FtsH plays a major role in the D1 degradation process. In *Chlamydomonas*, FtsH is composed of two subunits FtsH1 and FtsH2 and the *ftsh1-1* mutation, changing a conserved arginine residue, strongly impairs oligomerization and proteolytic activity (Malnoë et al., 2014). This mutation restores photosynthesis in a mutant lacking heme *c*_i_ of Cyt*b*_6_*f* (Malnoë et al., 2011). In addition, it impairs degradation of D1 in high light, slowing down repair of PSII during photoinhibition and leading to enhanced light-sensitivity in mixotrophic and photoautotrophic conditions (Malnoë et al., 2014). To address the question whether the FtsH protease is involved in the immediate post-translational degradation of D1 in the *ohp2* mutant, we crossed the original *10.1a* strain with a strain carrying the *ftsH1-1* mutation. In the progeny, the phenotypes associated with the WT, *ohp2*, *ohp2 ftsH1-1* and *ftsH-1* genotypes showed distinct fluorescence and growth patterns (Figure 6). When strains were analyzed by spot tests on agar plates, the *ftsH1-1* strains showed retarded growth and decreased Fv/Fm with increasing light intensity under photoautotrophic as well as under mixotrophic conditions (Figure 6A). This is typical of the *ftsH1-1* light-sensitive phenotype (Malnoë et al., 2014). As expected, the *ohp2* progeny did not grow photoautotrophically, except for occasional suppressor clones. Under mixotrophic conditions, they showed no deleterious effect of high light for growth, as is typical of PSII mutants, and had practically no Fv/Fm (Figures 6A, 6B). The *ohp2 ftsH1-1* strains showed interesting phenotypes. Like the *ohp2* parent, they were unable to grow on minimal medium at any light intensity tested (Figure 6A). However, their fluorescence induction curves were clearly indicative of the presence of a small amount of PSII, especially in dark-grown cells (Figure 6C). On TAP, we did not observe any stimulatory effect of low light on growth (Figure 6A). These results suggest that FtsH is involved in the degradation of the PSII units, possibly abnormal, produced in *ohp2* mutants. However, the activity of the low amount of PSII stabilized by attenuation of FtsH is either insufficient or too light-sensitive to allow photoautotrophy. Note that the unknown suppressor locus was not genetically linked to *FTSH1* in crosses, and the suppressor strains showed no sequence change in any of the *FTSH* genes, nor in any other known chloroplast protease gene.

**Figure 6.**
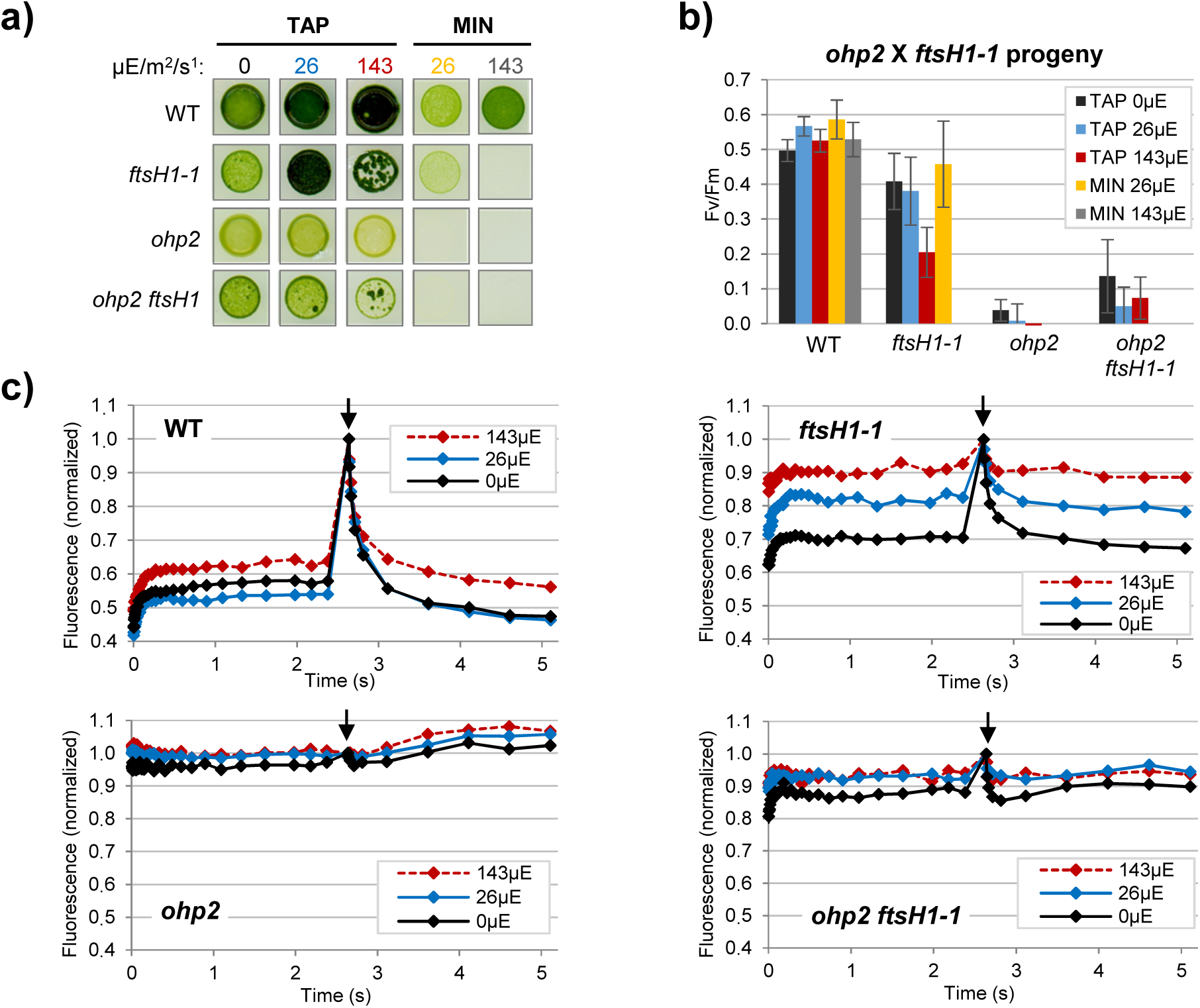
Mutation of the FtsH protease partially restores PSII activity but not photoautotrophy. a) Growth tests of the progeny of a cross *10.1a* X *ftsH1-1*. Cells were grown on TAP or MIN medium at the indicated light intensity. Spots shown are typical of the indicated genotypes. b) Fv/Fm values for the four genotypes, recorded from the plates in a). Values are average of 8, 7, 11 and 6 strains for the WT, *ftsH1*, *ohp2* and *ohp2 ftsH1* genotypes, respectively. Error bars represent S.D. c) Typical fluorescence induction curves for the four genotypes, recorded with the fluorescence camera on the TAP plates described in a). Fluorescence is normalized to the Fm value (arrow indicates saturating flash).

## DISCUSSION

### In *Chlamydomonas*, OHP2 is not necessary for *psbA* translation

The molecular analysis of an *ohp2* knockout mutant from *Chlamydomonas* revealed a major defect in PSII biogenesis, as indicated by its inability to grow photoautotrophically, the complete loss of PSII activity and the absence of the major PSII subunits, in particular D1 (Figures 1, 3). No effect was observed on the PSI RC (Figures 1C, 3A, 3D, 5), as reported for cyanobacterial HLIPs (Komenda and Sobotka, 2016), but at slight variance with land plants, where reductions in PSI subunits and antenna proteins have been reported (Beck et al., 2017; Li et al., 2019; Myouga et al., 2018). In *Chlamydomonas*, we observed an increased accumulation of LHCI antenna proteins in the mutant (Supplemental Dataset 2), and a higher abundance or stability of the PSI-LHCI complex in non-denaturing PAGE (Figure 3D). Whether this is related to the observed decrease in Chl abundance and changes in expression of Chl pathway genes remains to be investigated. In the Arabidopsis *ohp1* and *ohp2* mutants, the level of PSI RC decreases more than that of LHCI, but this appears to be true for other PSII mutants as well (Li et al., 2019)..

In other organisms, a dimer of One-Helix Proteins (HliC/D in cyanobacteria, OHP1/2 in land plants) has been found to mediate the early association of Chl *a* to the nascent D1 polypeptide. In *Chlamydomonas*, the expression patterns of OHP2, OHP1 and HCF244 in a variety of conditions are highly similar (Supplemental Figure S11) and on the Phytozome website (https://phytozome-next.jgi.doe.gov/) the latter two appear in each other’s lists of best correlated genes, pointing to the existence of a similar complex in the alga. We will therefore assume that the *Chlamydomonas ohp2* mutant lacks the OHC, which prevents normal cofactor insertion into D1, resulting in destabilization of PSII. Note that, although Chl is usually mentioned as the essential cofactor missing in *ohp2*, Phe *a* is also a possible substrate for the action of the OHC. The main result of our study, compared with those in higher plants, is that in *Chlamydomonas* the absence of OHP2 only marginally affects translation of the *psbA* mRNA. While our ^14^C pulse labelling experiments do point to the primary phenotype being a defect in D1 production, they can be interpreted either as a deficiency in *psbA* translation, or as a very fast degradation of the newly synthesized D1 polypeptide (Figure 3C). Our ribosome profiling experiments strongly support the latter hypothesis, as they show an almost normal abundance of FPs over the CDS (1.6-fold reduction of the signal). Continued translation of the *psbA* mRNA is observed over its whole length (Figure 4C), further supported by the fact that the proteomic analysis identified several peptides in *ohp2* that lie in the C-terminal part of D1.

Note that the two techniques we have used have their limitations. Usually, the absence of a signal in ^14^C-pulse labeling in *Chlamydomonas* is interpreted as loss of translation (e.g. de Vitry et al., 1989). But the *ohp2* mutant must be analyzed differently, as the mutation is supposed to affect a co-translational or early post-translational step. In addition, the sensitivity of this approach is limited by the broadness of the D1 band (D originally stands for “diffuse”) and the unavoidable presence of background radioactivity. Ribosome footprinting is more sensitive, even though it also is an indirect proxy for translation initiation. The abundance of RFs over a transcript may also be affected by the dynamics of translation, for example the stalling of ribosomes will increase the probability to generate RFs at this position. It is interesting to note that the reduction in ribosome occupancy in the *ohp2* mutant is more pronounced over the first half of the mRNA. Chl attachment to D1 has been proposed in barley to be associated with ribosome pausing (Kim et al., 1994a; Kim et al., 1991, 1994b) even though RF experiments in maize have failed to identify such Chl supply dependent pauses (Zoschke and Barkan, 2015; Zoschke et al., 2017). If interaction of the nascent chain with the Chl attachment machinery slows down elongation, then its absence in *ohp2* may reduce the density of RFs. Conservatively, we propose that in the absence of OHP2, *psbA* translation is maintained at a significant rate, over the whole CDS. However, the degradation of apo-D1 is so fast that full-length D1 remains below detection in pulse labeling experiments, as it is in Western blots (Figures 3A, 3C). This statement is backed by the observation that mutations in the FtsH protease and at an unknown suppressor locus can partially or fully restore PSII accumulation, which would be difficult to achieve if OHP2 was required for translation.

The situation is different in land plants, where OHP1, OHP2 and HCF244 are required for the recruitment of ribosomes to the *psbA* mRNA (Chotewutmontri et al., 2020). The abundance of RFs in the *Arabidopsis ohp2-1* knockout mutant was ~12 times lower than in the WT, a figure probably underestimated by normalization since the *psbA* CDS itself contributes to a large fraction of the RFs in the WT. Chotewutmontri et al. convincingly explained the discrepancy with a previous study using polysome analysis (Li et al., 2019), by the difficulty in analyzing *psbA* polysomes profiles due to the high abundance of the mRNA. They found that the *Arabidopsis ohp1-1* and maize *hcf244-1/-3* null mutants also fail to translate D1 (6-fold and 10-fold reduction, respectively). No change was observed in the pattern of RFs over *psbA* in any of the mutants, in line with a defect in translation initiation rather than elongation. All this was in accordance with the role of HCF244 as an essential translation initiation factor for *psbA* and its total absence in mutants lacking OHP1 or OHP2 (Li et al., 2019). Chotewutmontri and coworkers recently proposed a model for regulation of *psbA* translation by light, whereby the D1-bound OHC inhibits the ability of the stroma-exposed components (HCF244 and/or OHP2’s stromal tail) to initiate *psbA* translation (Chotewutmontri and Barkan, 2018; 2020; Chotewutmontri et al., 2020). It must be noted that Chotewutmontri et al. (2020) quantified RFs by Illumina read counting, while we have used hybridization to oligonucleotide probes. But both methods are fully validated, and the results are so different that we cannot invoke a technical bias. Therefore, we conclude that *Chlamydomonas* lacks the OHP2-dependent control mechanism for *psbA* translation described in land plants.

We used proteomics analyses to explain this apparent discrepancy in the context of the high conservation of the proteins between algae and land plants (as exemplified by our observation that *Arabidopsis* OHP2 can complement the *Chlamydomonas* mutant; Supplemental Figure S10). Quantification of OHP1 was difficult because of its small size, but the results of our targeted analysis point to the presence of traces of this protein in the *ohp2* mutant (Figure 5C). This, however, should have no effect on the insertion of cofactors: an *ohp2* mutation in *Arabidopsis* is not complemented by overexpression of OHP1 (Beck et al., 2017) and OHP1 alone does not bind pigments *in vitro* (Hey and Grimm, 2020). Studies in land plants have found that while some OHP2 can accumulate in *ohp1* null mutants, OHP1 is usually undetectable in *ohp2* mutants (Beck et al., 2017; Li et al., 2019). Some variability is seen in these studies, possibly due to the fact that the T-DNA insertion in *OHP2* lies within an intron (Beck et al., 2017; Li et al., 2019; Myouga et al., 2018). Similarly, the level of HCF244 reported in *Arabidopsis ohp2* mutants varied from nil to very low (Li et al., 2019) and the accumulation of HCF244, even when overexpressed, was found to be limited by that of OHP2 (Hey and Grimm, 2020; Li et al., 2019). Here, our untargeted and targeted LC-MS/MS analyses concur to demonstrate not only that HCF244 accumulates to significant levels in the *Chlamydomonas ohp2* mutant (~25% of WT), but that it can interact with the membrane (Figure 5C). This might be explained by the remaining traces of OHP1 acting as an anchor for HCF244 and stabilizing it, or to an intrinsically higher stability of *Chlamydomonas* HCF244 when not assembled. We propose that these remaining 25% HCF244 in the *Chlamydomonas ohp2* mutant are responsible for the maintenance of *psbA* translation at an appreciable rate. This does not exclude that an autoregulatory circuit exists in WT *Chlamydomonas* similar to that described by Chotewutmontri and Barkan (2020) in land plants. In this case, the absence of OHP1/OHP2 in the mutant may prevent the remaining HCF244 from sensing the presence of unassembled D1, leading to enhanced activation of *psbA* translation. In any event, the strong coupling observed in plants between Chl insertion and *psbA* translation initiation appears largely broken in *Chlamydomonas*. This coupling may be ancient in oxygenic photosynthesis, as judged from the existence of a similar complex in cyanobacteria. It might be interesting to examine whether the cyanobacterial complex also controls *psbA* translation. A major difference between *Arabidopsis* and *Chlamydomonas* is that Chl production depends entirely on light in the former, while the latter can produce it in the dark. Thus, the availability of Chl for integration into PSII could be used by Angiosperms to sense light, while algae (and cyanobacteria) would have to use other clues. In addition, the translation of *psbA* appears to mobilize a larger fraction of ribosomes in the plant than in the alga, as judged from RF and pulse-labeling experiments. Arguably, the alga can partly dispense of a regulatory circuit aimed at ensuring that *psbA* is translated only in the light. Detailed studies in other systems, such as land plants capable of producing Chl in the dark, are necessary to decide how strong the coupling needs to be to prevent deleterious effects of uncoordinated D1 synthesis and assembly, in particular during high light stress when D1 needs to be repaired at high rate.

### The absence of OHP2 affects translation of several chloroplast transcripts encoding PSII subunits

Interestingly, the *ohp2* mutant revealed large changes in the ribosome profiling pattern of PSII transcripts other than *psbA.* Most striking was the reduced translational output for the CES subunit CP47, encoded by *psbB* (Figure 4). This observation is in line with previous knowledge on translational regulation by assembly (the CES process) in *Chlamydomonas* (reviewed in Choquet and Wollman, 2009; Minai et al., 2006). Genetic studies indeed have demonstrated that in the absence of D1, the unassembled CP47 subunit feeds back onto the translation initiation of its own mRNA. Here, assuming that no assembly of CP47 and D1 can occur in the *ohp2* mutant, we can quantify this effect. With a 3-fold reduction of RFs over the *psbB* mRNA in the mutant (log_2_ *ohp2*/WT = −1.58), the CES effect appears to be rather dramatic (Figure 4B). Remarkably, the translation of *psbH* is also reduced in the mutant, in line with a complete lack of detection of PsbH peptides by proteomic analyses (Supplemental Datasets 2, 3). This supports the identification of PsbH as a component of the PSII CES cascade, as proposed recently by Trösch et al. (2018; Figure 4D). Early reports in *Chlamydomonas* describe a role of PsbH in assembly and/or stabilization of PSII (O’Connor et al., 1998; Summer et al., 1997) but the protein’s function has not been studied in detail. In the cyanobacterium *Synechocystis sp.* PCC 6803, PsbH is associated with CP47 and facilitates D1 processing and incorporation into PSII (Komenda et al., 2005). Assuming a similar position of PsbH in the assembly pathway in *Chlamydomonas,* its CES-mediated downregulation in the absence of D1 is consistent with that of CP47. The mechanism remains to be determined: is this regulation dependent upon the accumulation of unassembled PsbH, unassembled CP47 or the CP47/PsbH sub-complex? Genetic studies using mutants lacking the CP47 or PsbH proteins will be necessary to address this question. In their study, Chotewutmontri et al. (2020) pointed to an increased, rather than decreased, translation of *psbB* in their *ohp1*, *ohp2*, *hcf244* and even in *hcf173* mutants. The significance of these opposite behaviors between *Chlamydomonas* and land plants remains to be explored. Noticeably, *psbH*, compared to other PSII subunits, also appeared as slightly increased in the RF abundancy plots of the *ohp2* and *hcf244* mutants in Chotewutmontri et al. (2020).

The comparable effect that we observed on *psbT* translation in our experiments (Figure 4) could be construed as evidence that this small polypeptide also belongs to the PSII CES cascade. To our knowledge, no study has examined the possibility that PsbT also is subject to translational regulation. To complicate the matter, RF-analysis of the PSII mutant *nac2-26*, which lacks the D2 protein, rather showed a slight increase in translation of *psbT* (Trösch et al., 2018). Because D1 itself (and hence all the genes it controls via CES) needs D2 for full translation, the putative role of PsbT in the CES cascade remains ambiguous. It should be noted that all three cistrons belong to the conserved *psbB-T-H* operon and that the tetratricopeptide repeat protein MBB1 is necessary for stabilization of the 5’ end of the *psbB-psbT* mRNA as well as the processing and translation of the co-transcribed *psbH* mRNA (Loizeau et al., 2014; Vaistij et al., 2000). No monocistronic transcript has been described for PsbT, so its CDS can only be translated from the *psbB-psbT* dicistronic mRNA (Cavaiuolo et al., 2017; Monod et al., 1992). It is striking that the three ORFs that depend on MBB1 for transcript stabilization or translation showed a strong reduction of ribosome occupancy in the *ohp2* mutant (Figure 4C). This suggests a role for MBB1 in the CES-regulation: it may work as a translation factor or recruit such a factor. Even if MBB1 does not bind directly upstream of *psbT*, the increased recruitment of the *psbB-psbT* transcript to ribosomes may enhance translation initiation on *psbT*. Unfortunately, MBB1 could not be quantitated in our proteomic analyses. The decrease in RF level observed over the *psbN* ORF which is positioned between *psbT* and *psbH* but in antisense orientation could also indirectly involve MBB1, if recruitment of the precursor *psbB-T-H* transcript to the polysomes affects the stability/translatability of the antisense *psbN* mRNA. Coupling between chloroplast mRNA stabilization and translation factors has been reported in *Chlamydomonas*, for the MCA1/TCA1 couple. Changes in the abundance of MCA1/TCA1 rapidly regulate *petA* mRNA accumulation and translation to meet the cell’s demand for synthesis of the CES-regulated Cyt*f* subunit of the Cyt*b_6_f* complex (Loiselay et al., 2008; Raynaud et al., 2007). Similarly, the abundance of TAA1, involved in stabilization and translation of the *psaA* mRNA, has been reported to be involved in the downregulation of PSI subunits upon iron deficiency (Lefebvre-Legendre et al., 2015).

An unexpected result for the *ohp2* mutant in the RF experiment was the 2.8-fold increase in RFs over *psbD*, encoding the main RC subunit D2 (Figures 4A, 4B; Supplemental Figures S6B, S7; Supplemental Dataset 1). Because of the high variability in the efficiency of ^14^C incorporation between strains and of the gel background, we could not verify this by the pulse-labeling experiments (Figure 3C). However, as all other D1 mutants studied thus far, like FuD7 or F35, completely lacked D1 translation (Bennoun et al., 1986; Girard-Bascou et al., 1992; Yohn et al., 1996), a stimulatory effect on D2 translation by the presence of unassembled D1 (as possible in *ohp2*) could not have been observed. We note that translation of *psbD* was not enhanced in any of the mutants studied by Chotewutmontri et al. (2020).

### OHP2 is not completely essential for PSII biogenesis

Another intriguing characteristic of the *Chlamydomonas ohp2* mutant is the high frequency at which the non-photoautotrophic phenotype can be suppressed. A mere plating on mineral medium, without any mutagenic treatment, was enough to generate dozens of extragenic suppressors. In spite of all our efforts, we were unable to identify a plausible causative variant, even though genetic analysis of three independent suppressors indicated that the suppressor mutation(s) were tightly linked, but unlinked to the *OHP2* or *FTSH1* loci. Furthermore, combination of *ohp2* with the *ftsH1-1* mutation that severely affects activity of the FtsH protease (Malnoë et al., 2014) allowed partial recovery of stable PSII charge separation, as revealed by measurements of variable fluorescence and ECS (Figure 6C). This indicates that whatever the exact action of the OHC on D1, it is not completely essential for biogenesis of a functional PSII. A by-pass reaction must be possible, that leads to partial restoration of PSII biogenesis. This pathway would be activated by the suppressor mutation, but it probably already exists in the *ohp2* mutant: by slowing down D1 degradation, the *ftsH1-1* mutation may allow a normally unstable form of D1 to proceed through this alternative pathway. Such a pathway also operates in cyanobacteria, where deletion of all HLIP genes does not prevent PSII biogenesis (Xu et al., 2004).

It is probable that the PSII produced by the OHP2-independent pathway is not fully functional, because the suppressed strains showed some light-sensitivity, and the *ohp2 ftsH1-1* double mutant was not photoautotrophic (Figure 6A). Alternatively, the activity of this alternative pathway may be too low to allow efficient repair in high light. The *ftsH1-1* mutation by itself renders the cell highly sensitive to photoinhibition. Detailed studies comparing PSII purified from these strains may be necessary to answer this question.

### Consequences of the absence of OHP2 on other steps of the PSII assembly pathway

Based on the above, it was interesting to characterize the assembly intermediates in the mutant and suppressor. We therefore used our proteomics data to monitor the accumulation of the many PSII assembly factors described in the literature. Not only did they represent candidate genes for the suppressor mutation, but changes in their accumulation in the mutant and suppressor could reveal regulatory mechanisms at play in *ohp2* and the suppressor. Some of the changes described below could be brought about by regulation of gene expression, others by (de)stabilization of the proteins. By comparing whole cells and membranes, we may also detect changes in the association of these factors with PSII assembly intermediates, as recently demonstrated by Spaniol et al. (2021) for the *lpa2* mutant. Figure 7 proposes a working model for the role of the OHC and associated proteins in D1 synthesis, combining previous reports from other organisms with data presented here for *Chlamydomonas*. In the WT (Figure 7, left panel), HCF173 binds the 5′-UTR of *psbA* mRNA to promote its translation (Link et al., 2012; Schult et al., 2007; Williams-Carrier et al., 2019), which is then likely triggered by HCF244. Nascent pD1 protein is co-translationally inserted into the thylakoid membrane where it is stabilized by membrane-associated luminal HCF136. The OHC is proposed to further stabilize pD1 by the insertion of Chl/Phe/Car, allowing its assembly in the RC complex and formation of functional PSII core, dimer and finally supercomplexes. A striking result of our targeted proteomics analysis was the two-fold overaccumulation of HCF173 in the membranes of the *ohp2* mutant (Figure 5C, Supplemental Dataset 4). It was not significantly overexpressed in the mutant at the whole cell level, so its increased abundance in membranes probably indicates that a larger fraction of the *psbA* mRNA is undergoing translation and thus found on membrane-associated polysomes. This effect is no longer seen in the suppressor, suggesting that it is not a direct consequence of the absence of the OHC, but rather a compensatory mechanism caused by the PSII deficiency. As proposed above, unassembled D1 can no longer regulate the activity of HCF244 in the mutant because the OHC cannot form (Chotewutmontri et al., 2020; Link et al., 2012). This is expected to lead to enhanced translation initiation on the *psbA* mRNA driven by the residual but fully active HCF244 (Figure 7, middle panel), unless a counteracting mechanism signals that PSII biogenesis proceeds, as in the suppressor. SRRP1, another known interactant of HCF173 and of the *psbA* mRNA, is believed to act as a repressor of translation (Watkins et al., 2020). It could not be quantitated in the membranes, but interestingly, while abundant in cells of the mutant and WT it was undetectable in the suppressor, even though the gene appeared unaltered.

**Figure 7.**
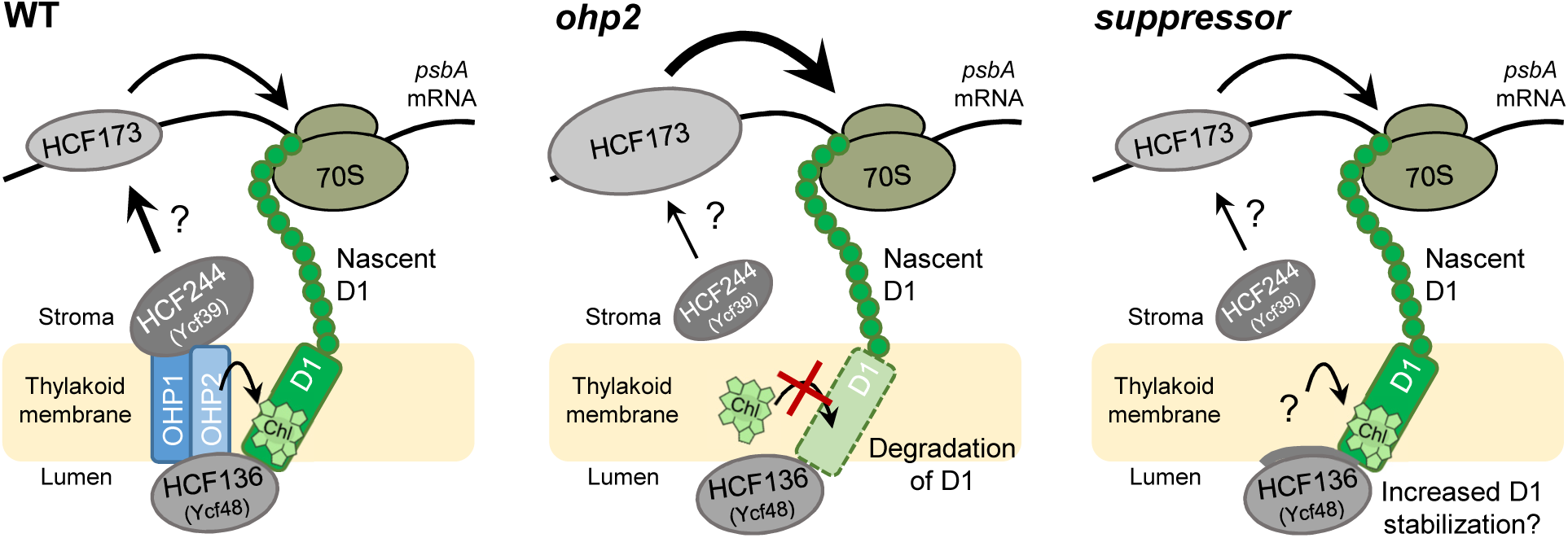
Schematic model for the role the OHC complex and associated proteins in D1 synthesis and first steps of PSII de novo assembly. In the WT (left panel), likely triggered by HCF244, HCF173 binds the 5′-UTR of *psbA* mRNA to promote translation initiation. pD1 is co-translationally inserted into the thylakoid membrane. Membrane associated luminal HCF136 stabilizes pD1 and is involved in RC assembly. The OHC is required for early PSII assembly steps and proposed to insert Chl into pD1 proteins. Recently reported negative autoregulatory circuits of *psbA* translation initiation involving the OHC are not displayed here (Chotewutmontri & Barkan, 2020; Chotewutmontri et al., 2020). In *ohp2* (middle panel) and suppressor strains (right panel) residual levels of HCF244 may promote *psbA* translation via interaction with HCF173. Ongoing *psbA* translation may be further supported by overaccumulation of HCF173. However, in the *ohp2* mutant synthesized D1 is rapidly degraded and does not accumulate in the absence of the OHC complex. In the suppressor strains, stabilization of nascent D1 may be accomplished by increased membrane association of HCF136 or other mechanisms to allow assembly of early PSII intermediates. HCF173, HCF136, and HCF244 protein amounts detected by proteomics are indicated by differently sized ovals. Names of cyanobacterial homologs are given in brackets.

HCF136 is a conserved luminal protein, known in cyanobacteria as Ycf48. Its role seems mostly to promote the association of the pD1/PsbI subcomplex with a distinct precomplex consisting of D2 and the heterodimeric Cyt*b_559_* (formed by the PsbE and PsbF subunits), to form the RC (reviewed in Nickelsen and Rengstl, 2013; Zhang et al., 1999). It is required for the assembly of early PSII intermediates, possibly with the concurrent incorporation of Chl and has been found to bind pD1 (but not mature D1) in a split-ubiquitin assay (Hey and Grimm, 2020; Knoppová et al., 2014; Komenda et al., 2008; Li et al., 2019; Meurer et al., 1998; Myouga et al., 2018; Plücken et al., 2002). It was therefore striking to observe that association of HCF136 with the membrane (not its overall abundance) was markedly increased in the suppressor, compared to the WT or mutant (Figure 5; Supplemental Datasets 2-4). While genome sequencing rules out *HCF136* as being the suppressor locus, its increased abundance in the membranes of the suppressor may indicate the stabilization of a complex comprising pD1, allowing it to associate with Chl via the alternative pathway proposed above (Figure 7, right panel).

Assembly factors acting downstream of the formation of the RC can also be impacted by the absence of OHP2. For example, the mutant did not show any association of PSB28 with the membrane, consistent with the fact that this protein binds to the RC47 complex (Zabret et al., 2021), that cannot form in the mutant. This effect was reverted in the suppressor, in line with a restoration of the assembly pathway. PSB27 binds to CP43 on the luminal surface of the PSII monomer and dimer and contributes to the formation of the OEC (Avramov et al., 2020; Huang et al., 2021; Zabret et al., 2021). In the *ohp2* mutant, it overaccumulates at the whole-cell level, presumably through enhanced expression of the gene, but not in the membranes, probably because of the absence of the RC. In contrast, the suppressor showed enhanced binding of PSB27 to the membrane, suggesting that PSII formed in the absence of the OHC requires prolonged association with PSB27, for example because pD1 maturation, binding of the OEEs and/or photoactivation are delayed. Another interesting observation that links back to the role of FtsH is the increased association of PSB29/THF1 with the membrane in *ohp2*. The *Arabidopsis thf1* mutant is variegated (Wang et al., 2004), a phenotype suppressed by mutations affecting chloroplast gene expression (Hu et al., 2015; Zhang et al., 2009), just like that of the *ftsH* mutants. THF1 is necessary for normal accumulation and function of FtsH in land plants and cyanobacteria (Becčková et al., 2017; Huang et al., 2013; Zhan et al., 2016). Its 3D structure has been solved (Becčková et al., 2017), but it is unclear yet how it stabilizes or activates the protease. In our proteomic analysis, PSB29/THF1 was not over accumulated in any of the strains. However, its increased association with the membrane in the mutant may reflect the increased activity of FtsH involved in degrading ill-conformed D1.

### Other effects of the mutation in *OHP2* on chloroplast biogenesis

Our RF experiments showed an increased translation of all the chloroplast-encoded subunits of the ATP synthase in the *ohp2* mutant (Supplemental Figure S6, Supplemental Dataset 1). Similarly, our proteomics experiments revealed an increased abundance of all subunits of the complex (including the nucleus-encoded ATPC/D/G), with a striking quantitative coherence in membrane samples (Figures 5A, 5B; Supplemental Datasets 2, 3). Increased levels of chloroplast ATP synthase subunits were recently also observed in complexome profiling of the PSII-deficient *lpa2* mutant from *Chlamydomonas* (Spaniol et al., 2021). Overaccumulation of the ATP synthase could be involved in maintaining energy balance in the chloroplast: in the absence of PSII, linear electron flow is abolished, but cyclic electron flow remains possible between PSI and the Cyt*b*_6_*f* complex and generates a proton gradient that can be used for ATP synthesis.

Our proteomics analysis showed increased accumulation of LHCI subunits in the *ohp2* mutant, as well as in the suppressor strain. This suggests that the lack of OHP2 directly affects the biogenesis of PSI. Interestingly, BN-PAGE revealed a stronger association of the PSI core and antenna subcomplexes in the mutant. This may be related to the observation that the PSAN subunit, although not affected in whole cells, was undetectable in the membranes of the mutant. In maize, PSAN lies at the interface between the RC and LHCI (Pan et al., 2018), and this subunit was not found in the structure of the Chlamydomonas PSI-LHCI complex (Suga et al., 2019), suggesting that it modulates the association between the antenna and the RC in PSI.

Proteomics also revealed a general stimulation of the heme/Chl biosynthesis pathway in the *ohp2* mutant (Figures 5A, 5B; Supplemental Datasets 2, 3). To our knowledge, this is not a general feature of PSII deficient mutants and may be specifically linked to the main purported function of OHP2/OHP1, i.e. integration of Chl into a nascent polypeptide. Accumulation of the protochlorophyllide *a* oxidoreductase POR1 was increased both at the cell and membrane level, while for other enzymes like the Mg-chelatase, the Mg-protoporphyrin-IX methyltransferase CHLM and the cyclase CTH1, the observed effect was an increased association with the membrane. The OHC complex has been linked before with Chl synthesis, albeit in different manners. In cyanobacteria, the Chl synthase ChlG (found unaffected here in the *Chlamydomonas ohp2* mutant) co-immunoprecipitates with the homologs of OHP2 and HCF244 (Chidgey et al., 2014). In the *Arabidopsis ohp2* mutant, Hey and Grimm (2018) reported a decreased accumulation for most of the immunologically analyzed Chl biosynthesis enzymes. The authors proposed a posttranslational destabilization of these proteins in response to OHP2 deficiency, which could be beneficial when D1, a major sink for newly synthesized Chl, is not translated. In *Chlamydomonas*, the continued translation of D1, when the absence of OHP2 limits its ability to ligate Chl, may instead start a signaling cascade aimed at increasing production of Chl. The *ohp2* mutant may be missing a negative feedback loop normally operating in the WT. For example, a product of the interaction between the OHC and pD1, such as the Chl-associated but unassembled pD1, could negatively regulate association of the Chl synthesis enzymes to the membrane, signaling that more pD1 is made than can be assembled and the Chl flux should be lowered. Similarly, the activity of HCF244 as a translation activator has been proposed to be regulated by the interaction of pD1 with the OHC in the dark (Chotewutmontri and Barkan, 2020). Another striking observation is the marked Chl deficiency in the mutant (Supplemental Table S1). The PSII core contains 35 Chl *a* molecules, while its pigment bed harbors ~200 Chl, so loss of the core itself cannot fully account for the 40% decrease in Chl content that we observed. In the *Arabidopsis ohp2* mutants, a comparable or even stronger reduction in Chl accumulation was observed, with Chl *b* reduced to a lesser extent than Chl *a* (Hey and Grimm, 2018; Myouga et al., 2018). A regulatory role of the OHC on Chl synthesis might be worth further exploration.

## MATERIAL AND METHODS

### Strains and culture conditions

The *ohp2* mutant (*10.1.a*) was isolated in a screen and was generated by nuclear transformation of the mating type minus (mt-) cell-walled recipient strain Jex4 with the plasmid pBC1 cut with SacI and KpnI, described in Houille-Vernes et al. (2011). For backcrossing of *ohp2* strains, the wild type WT-S34 was used. For localization studies the cell wall deficient *UVM4* strain generated by Neupert et al. (2009) was used as recipient strain for glass bead transformation. Two PSII mutants, *nac2-26* which lacks the *psbD* mRNA stabilization factor NAC2, and the *psbA* deletion mutant FuD7, served as controls (Bennoun et al., 1986; Boudreau et al., 2000).

Algal strains were grown at 23°C under continuous white light on Tris-acetate-phosphate medium (TAP) at ~10-20 µE.m^−2^.s^−1^, or on High Salt Minimum medium (MIN) agar plates at 100 µE.m^−2^.s^−1^, respectively (Harris, 2009). *ohp2* mutant strains were kept on plates at very low light (~ 5 µE/m^2^/s) to limit inadvertent selection of suppressor mutants. Before usage of *ohp2* in experiments the absence of potential PSII suppressors was confirmed by Fv/Fm measurements. For experiments, liquid cultures of *Chlamydomonas* were grown in TAP medium supplemented with 1% sorbitol (TAPS) at ~20 – 30 µE.m^−2^.s^−1^ until a cell density of 2-3 * 10^6^ cells/mL was reached.

The reversion rate of *ohp2,* was determined according to Kuras et al. (1997). *ohp2* mutant cells were collected by centrifugation and resuspended in HSM to a density of 2 × 10^8^ cells/mL. 0.5 mL was spread onto 10 HSM agar plates and maintained under continuous light for three weeks, at which time colonies were counted.

### Spectroscopy and fluorescence measurements

In vivo spectroscopy measurement using a JTS-10 spectrophotometer (Biologic, Grenoble, France) was performed as described before in Jalal et al. (2015), using cells grown under low light conditions. Fv/Fm is presented as the average of four fluorescence induction measurements at 26, 56, 135 and 340 µE/m^2^/s followed by a saturating pulse. PSII/PSI ratio was calculated from the difference in ECS signal at 520 nm, measured 160 µs after a saturating single turnover flash, in the absence or presence of DCMU (10 µM) and hydroxylamine (1 mM) to measure (PSI+PSII) or PSI activity, respectively. In the latter condition, the b/a ratio is the ratio of amplitude of the second phase of the ECS signal (phase b, due to proton pumping by Cyt*b*_6_*f*) to the initial phase a (PSI) and is a rough measure of Cyt*b*_6_*f* activity.

### Analysis of nucleic acids

For isolation of nucleic acids, cells were harvested by centrifugation at 1,100 × g, 4°C for 6 min. Genomic DNA was extracted using the DNeasy Plant Mini Kit (Qiagen, Hilden, Germany) according to manufacturer’s protocol or CTAB buffer (2% cetyltrimethylammonium bromide, 100 mM Tris-HCl, pH 8) followed by phenol/chloroform/isoamyl alcohol extraction (25:24:1, Carl Roth GmbH, Mannheim, Germany). DNAs were digested by restriction enzymes and separated on 0.8% agarose gels set up in TPE-buffer (89 mM Tris-phosphate, 2 mM Na_2_EDTA). Total cellular RNA was extracted by using the TRI reagent (Sigma-Aldrich, Saint Louis, USA), according to the manufacturer’s instructions. RNAs were separated on 1% denaturing formaldehyde agarose gels. After separation nucleic acids were transferred to Roti Nylon^+^ membrane (Roth, Karlsruhe, Germany), followed by UV light cross-linking (UV Crosslinker, UVC 500, Hoefer Inc., San Francisco, USA). Dig-labeled probes were synthesized by PCR from total DNA or cloned cDNA (*psbA*) by using primers denoted in Supplemental Table S2. Hybridizations and detection of dig-labeled probes were performed using standard methods.

### Immunoblot analysis

For isolation of total protein extracts, cells from 20 mL liquid cultures were harvested by centrifugation and cell pellets resuspended in 200 µL 2 × lysis buffer (120 mM KCl, 20 mM tricine pH 7.8, 5 mM β-mercaptoethanol, 0.4 mM EDTA, 0.2% Triton X100) supplemented with protease inhibitors (cOmplete^™^ ULTRA Tablets, Mini, Roche, Switzerland) and lysed via sonication on ice. To separate membrane proteins from soluble proteins for cell subfractionation, cells from 50 mL culture were resuspended in 500 µL hypotonic solution (10 mM Tricine/KOH pH 6.8, 10 mM EDTA, 5 mM β-mercaptoethanol, and protease inhibitors). Lysis was performed mechanically by vortexing thoroughly with glass beads (0.5 mm diameter) two times for 1 min. After centrifugation at 15,000 g for 10 min, the supernatant was considered as total soluble protein extract. The pellet containing the membrane proteins was resuspended in 2 × lysis buffer.

Protein concentration was measured by Bradford Protein Assay (Bradford, 1976). SDS-PAGE, protein gel blotting and immunodetection were performed as described by Sambrook and Russel (2001). Antibodies were as follows: α-PsaA (Agrisera, #AS06 172), α-D1 (Agrisera, #AS05084), α-CP43 (Agrisera, #AS11 1787) α-Cyt*f* (Agrisera, # AS08 306), α-HA (Sigma-Aldrich, #H6908). The antiserum against the spinach Rubisco holoenzyme used for the detection of RbcL was kindly provided by G. F. Wildner (Ruhr-University Bochum). The antibody against the *Chlamydomonas* D2 protein was generated by immunization of rabbits using recombinant D2-GST protein (BioGenes GmbH, Berlin, Germany).

### Complementation and localization studies

Constructs generated for the complementation of the *ohp2* mutant and OHP2 localization studies are shown in Supplemental Figure S3. All constructs were created by the insertion of a PCR-amplified sequence of interest into the pBC1-CrGFP vector (= pJR38, Neupert et al., 2009; construct 1, Supplemental Figure S3) via NdeI or NdeI/EcoRI restriction sites. Primers used for cloning are given in Supplemental Table S2. For cloning details of the synthetic *Arabidopsis* gene (construct 6, Supplemental Figure S3) please see Supplemental Data S2. For localization studies, the cell wall deficient strain *UVM4* was transformed by the glass bead method (Kindle, 1990; Neupert et al., 2009) and positive transformants were selected by growth on TAP plates supplemented with 10 µg/mL paromomycin (Pm). For complementation, constructs were integrated into the genome of the *ohp2* mutant strain by the electroporation method (Shimogawara et al., 1998), selected for Pm resistance and subsequently screened for photoautotrophic growth on HSM plates.

### 2D blue native (BN) PAGE

500 mL liquid cultures of *Chlamydomonas* strains were harvested at 1,000 × g for 10 min at 4°C and resuspended in 1 mL TMK buffer (10 mM Tris/HCl, pH 6.8, 10 mM MgCl_2_, 20 mM KCl) and protease inhibitors (cOmplete^™^ ULTRA Tablets, Mini, Roche, Switzerland). Resuspended cells were lysed by sonication (ultrasound pulses of 10 sec for 3 times). Cells were centrifuged (1,000 × g, 1 min, 4°C) to remove cell debris and unlysed cells. The supernatant was centrifuged for 10 min at 20,000 × g at 4°C. The resulting pellet was washed twice with TMK buffer and finally resuspended in 500 µL TMK buffer. Chl concentration was determined by adding 20 µL of the sample to 980 µL methanol, 5 min incubation at RT and a 1 min centrifugation step to remove the starch. OD652 of the supernatant was measured and Chl concentration calculated using the formula: Chl *a* (mg/mL) = A652nm * 1.45. Aliquots containing 25 μg of Chl were again centrifuged for 10 min, 20.000 g at 4°C and resuspended in 51 µL ACA buffer (750 mM ε-aminocaproic acid, 50 mM Bis-Tris pH 7.0, 5 mM pH 7.0 EDTA, 50 mM NaCl) and solubilized for 10 min on ice after addition of n-dodecyl-ß-D-maltoside (ß-DM) to a final concentration of 1.5%. Solubilized proteins were separated from insoluble material by centrifugation for 20 min, 16,000 × g at 4°C and mixed with 1/10 volume of BN loading dye (750 mM ε-aminocaproic acid, 5% Coomassie G250 (w/v)). First dimension native electrophoresis was carried out according to Schägger and Jagow (1991) on a 5 – 12 % linear gradient. Protein complexes were further subjected to 2D 12% SDS-PAGE and analyzed by immunoblotting.

### Ribosome profiling and data analysis

For each replicate, 500 mL culture was supplemented with 100 µg/mL chloramphenicol and 100 µg/mL cycloheximide, followed by rapid cooling using plastic ice cubes. The cold (< 8°C) culture was then centrifuged at 3,000 *g* for 2 min to pellet the cells. The pellet was washed once with ice-cold polysome buffer (20 mM Tris pH 8.0, 25 mM KCl, 25 mM MgCl_2_, 1 mM dithiothreitol, 100 µg/mL chloramphenicol, 100 µg/mL cycloheximide) and then flash-frozen in liquid nitrogen. The pellet was resuspended in 8 mL of polysome buffer containing 1x Protease inhibitor Cocktail (Roche) and 1 mM PMSF, and the cells were lysed using a French press (Avestin®) at 2 bar. Polysome isolation from the lysate, RNA digestion and FP isolation have been performed as described in Trösch et al. (2018). For the total RNA, a separate 8 mL of the culture was centrifuged at 3000 *g* for 2 min, and 750 µL of Trizol was added to the pellet directly. Total RNA extraction as well as RNA labelling and microarray analysis was performed as described in Trösch et al. (2018).

Data processing and analysis was conducted according to previous studies (Trösch et al., 2018; Zoschke et al., 2013): Local background was subtracted from single channels (F635-B635 and F532-B532, respectively) and probes located within annotated and confirmed chloroplast reading frames were normalized to the average signal of the compared datasets including all replicates of ribosome FPs and total mRNA to remove overall differences introduced by technical variations. Probes covering a respective ORF were averaged, and relative abundance of ribosome FPs and total mRNA was calculated for each ORF by normalizing each ORF value to the average of all ORF values. By this, expression of the individual ORF is considered in relation to mean values of all plastid-encoded genes. Relative translation efficiency is determined by comparing to values of average FP intensities relative to the average RNA intensities, for each ORF. All average values and standard deviations are based on three independent biological replicates. Significant differences in gene-specific RNA and ribosome FP accumulation and translation efficiencies between WT and *ohp2* mutant data were determined with a Welch’s t-test and corrected for multiple testing according to Storey’s q-value method. Genes were marked as significant for q-value of <0.05 and with expression changes more than two-fold (Supplemental Dataset 1).

### Mass spectrometry

#### Whole cell proteomics

50 µg of proteins were separated by SDS-PAGE. After a short migration (< 0.5 cm) and Coomassie blue staining, gel bands containing proteins were excised and destained. Gel pieces were subjected to a 30 min reduction at 56°C and a 1 h cysteine alkylation at room temperature using 10 mM dithiothreitol and 50 mM iodoacetamide in 50 mM ammonium bicarbonate, respectively. Proteins were digested overnight at 37°C using 500 ng of trypsin (Trypsin Gold, Promega). Supernatants were kept and peptides remaining in gel pieces were further extracted with 1% (v/v) trifluoroacetic acid. Corresponding supernatants were pooled and dried. Peptide mixtures were subsequently reconstituted in 200 µL of solvent A (0.1% (v/v) formic acid in 3% (v/v) acetonitrile). Five microliters of peptide mixtures were analyzed in duplicate on a Q-Exactive Plus hybrid quadripole-orbitrap mass spectrometer (Thermo Fisher) as described in Pérez-Pérez et al. (2017) except that peptides were separated on a PepMap™ RSLC C18 Easy-Spray column (75 µm × 50 cm, 2 µm, 100 Å; Thermo Scientific) with a 90 min gradient (0 to 20% B solvent (0.1% (v/v) formic acid in acetonitrile) in 70 min and 20 to 37% B solvent in 20 min).

#### Membrane proteomics

Cells were collected by centrifugation, resuspended in 25 mM phosphate buffer at a density of app. 0.5 µg/µL total Chl and disrupted by three consecutive freeze and thaw cycles. Soluble proteins were separated by centrifugation at 4°C, 25,000 *g* for 15 min. 20 µg total protein from the pellet fraction were precipitated in 80 % acetone, tryptically digested and desalted as described (Hammel et al., 2018). Peptides were resuspended in a solution of 2% acetonitrile, 1% formic acid just before the LC-MS/MS run. The LC-MS/MS system (Eksigent nanoLC 425 coupled to a TripleTOF 6600, ABSciex) was operated basically as described for data dependent acquisition (Hammel et al., 2018).

For targeted MRM-HR data acquisition a list of 39 precursor m/z was chosen for targeting the 5 different proteins. The precursors were selected from results of the data dependent acquisitions and good responding peptides as predicted by a deep learning algorithm for peptide detectability prediction, d::pPop (Zimmer et al., 2018). Collision energies were calculated from the standard CE parameters of the instrument, dwell time was set to 60 ms, fragments were acquired from 110 m/z – 1600 m/z, resulting in a cycle time of 2.6 s.

#### Data processing and label-free quantification

Shotgun proteomics raw data were processed using the MaxQuant software package as described in Martins et al. (2020) with slight modifications. For protein identification and target decoy searches were performed using a home-made *Chlamydomonas reinhardtii* protein database consisting in the JGI Phytozome nuclear-encoded proteins database (v.5.6) concatenated with chloroplast- and mitochondria-encoded proteins in combination with the Maxquant contaminants.

For whole cell proteomics the mass tolerance in MS and MS/MS was set to 10 ppm and 20 mDa, respectively and proteins were validated if at least 2 unique peptides having a protein FDR < 0.01 were identified. For quantification, unique and razor peptides with a minimum ratio count ≥ 2 unique peptides were used and protein intensities were calculated by Delayed Normalization and Maximal Peptide Ratio Extraction (MaxLFQ) according to Cox et al. (2014). For membrane protein data dependent runs MaxQuant software (v1.6.0.1) (Cox and Mann, 2008; Tyanova et al., 2016) was used. Peptide identification, protein group assignment and quantification was done with standard settings for ABSciex Q-TOF data except that 3 miss-cleavages were allowed, minimum peptide length was set to 6 AA, maximum peptide mass 6600 Da.

*Shotgun data analysis:* For statistical analysis, LFQ intensities obtained from MaxQuant were further analyzed with the Perseus software package version 1.6.15.0 (Tyanova et al., 2016). In case of the whole cell data the two measurement replicates were averaged first. For both datasets the biological replicates were grouped and the LFQ intensities were Log_2_ transformed. The data were filtered to contain at least three valid values (out of four biological replicates) or two valid values (out of three biological replicates) in case of the whole cell or membrane fractions, respectively. Significant changes in LFQ intensities compared to the WT were identified by a modified two sample t-test of the Perseus software (permutation-based FDR=5%, artificial within group variance S0=1). The *ohp2* mutant, the suppressor mutant *M-su1* and in case of the whole cell samples the complemented line *ohp2:OHP2-HA,* were compared to the WT independently. Data visualizations shown in Figure 5 and Supplemental Figure S9 were done using R (R Core Team, 2018). The log_2_ transformed LFQ intensities and t-test results of the whole cell samples and membrane fractions are shown in Supplemental Dataset 2 and Supplemental Dataset 3, respectively.

*Targeted mass spectrometry data analysis* was done using Skyline software (v20.1.0.155) (MacLean et al., 2010; Pino et al., 2020). The spectral library was built by prediction using Prosit (Gessulat et al., 2019), retention times were taken, where possible, from identification results of the respective peptides using MaxQuant. A minimum of 2 precursors per protein and 4 transitions per precursor was used for quantification. The integration borders were manually adjusted where necessary and dotp values for the coeluting fragments as reported from Skyline was taken as a quality measure for correct assignment (Supplemental Dataset 4). Retention times varied over all runs by less than 0.5 min, dotp values were usually >0.85 across the replicates where a correct assignment was assumed. Summed fragment peak areas were normalized for unequal sample loading to the total intensities of proteins as identified by MaxQuant in data dependent runs.

The untargeted mass spectrometry proteomics data have been deposited to the ProteomeXchange Consortium via the PRIDE (Perez-Riverol et al., 2021) partner repository with the dataset identifier PXD031558 (Reviewer account details: https://www.ebi.ac.uk/pride/login; Username; Password: D3xVDFGU). The targeted mass spectrometry proteomics data have been deposited to the ProteomeXchange Consortium via the Panorama Public (Sharma et al., 2018) partner repository with the dataset identifier PXD031631 (Reviewer account details: https://panoramaweb.org/TIFMYs.url; Username; Password: iEqMdcrZ).

## AUTHOR CONTRIBUTIONS

AVB, OVA, FWi, JN, and MS conceived and designed the research; FWa, KD, IM, FE, RT, FS, OVA, AVB, and XJ performed the experiments; FS, LDW, and FWi analyzed data; AVB, OVA, and FWi wrote the paper with inputs from all authors.

## ACKNOWLEDGEMENTS

The authors would like to thank C. de Vitry for providing the *ftsH1-1* strain.

## SUPPLEMENTAL MATERIAL

## SUPPLEMENTAL FIGURES

**Figure S1.** Genetic analysis of the *10.1a* mutant.

**Figure S2.** Identification of the *ohp2* mutation.

**Figure S3.** DNA constructs used for complementation and localization studies.

**Figure S4.** Hydrophobicity prediction for *Chlamydomonas* and *Arabidopsis* OHP2 amino acid sequences.

**Figure S5.** OHP2 is a membrane localized chloroplast protein.

**Figure S6.** Reproducibility of RF experiments and results for all chloroplast encoding genes.

**Figure S7.** Targeted ribosome profiling of chloroplast translation reveals enhanced *psbD* translation in the *ohp2* mutant.

**Figure S8.** Re-accumulation of D2 and restoration of photoautotrophy in suppressor mutants.

**Figure S9.** Heat maps representing the reproducibility of LC-MS experiments and complementation of *ohp2*.

**Figure S10.** Complementation of the *ohp2* mutant with the orthologous *Arabidopsis* protein restores photoautotrophy and D1 accumulation.

**Figure S11.** Transcript accumulation level of *OHP2*, *OHP1*, and HCF244 in *Chlamydomonas* under various growth conditions.

## SUPPLEMENTAL TABLES AND DATA

**Supplemental Table S1.** Photosynthetic parameters and Chl composition of *Chlamydomonas* WT Jex4 (WT), the *ohp2* mutant and complemented strain.

**Supplemental Table S2.** Primers used in this study.

**Supplemental Data S1.** Flanking sequence tags in the *ohp2* mutant obtained by Illumina sequencing.

**Supplemental Data S2.** Synthetic *Arabidopsis OHP2* nucleotide sequences and derived protein sequence used to complement the *Chlamydomonas ohp2* mutant strain.

## SUPPLEMENTAL DATASETS

**Supplemental Dataset 1.** Complete dataset from array-based ribosome profiling experiments.

**Supplemental Dataset 2.** Complete dataset from whole cell proteomics experiments.

**Supplemental Dataset 3.** Complete dataset from membrane proteomics experiments.

**Supplemental Dataset 4.** Targeted mass spectrometry data

## Supplemental Tables, Data and References

### Supplemental Figures

**Figure S1.**
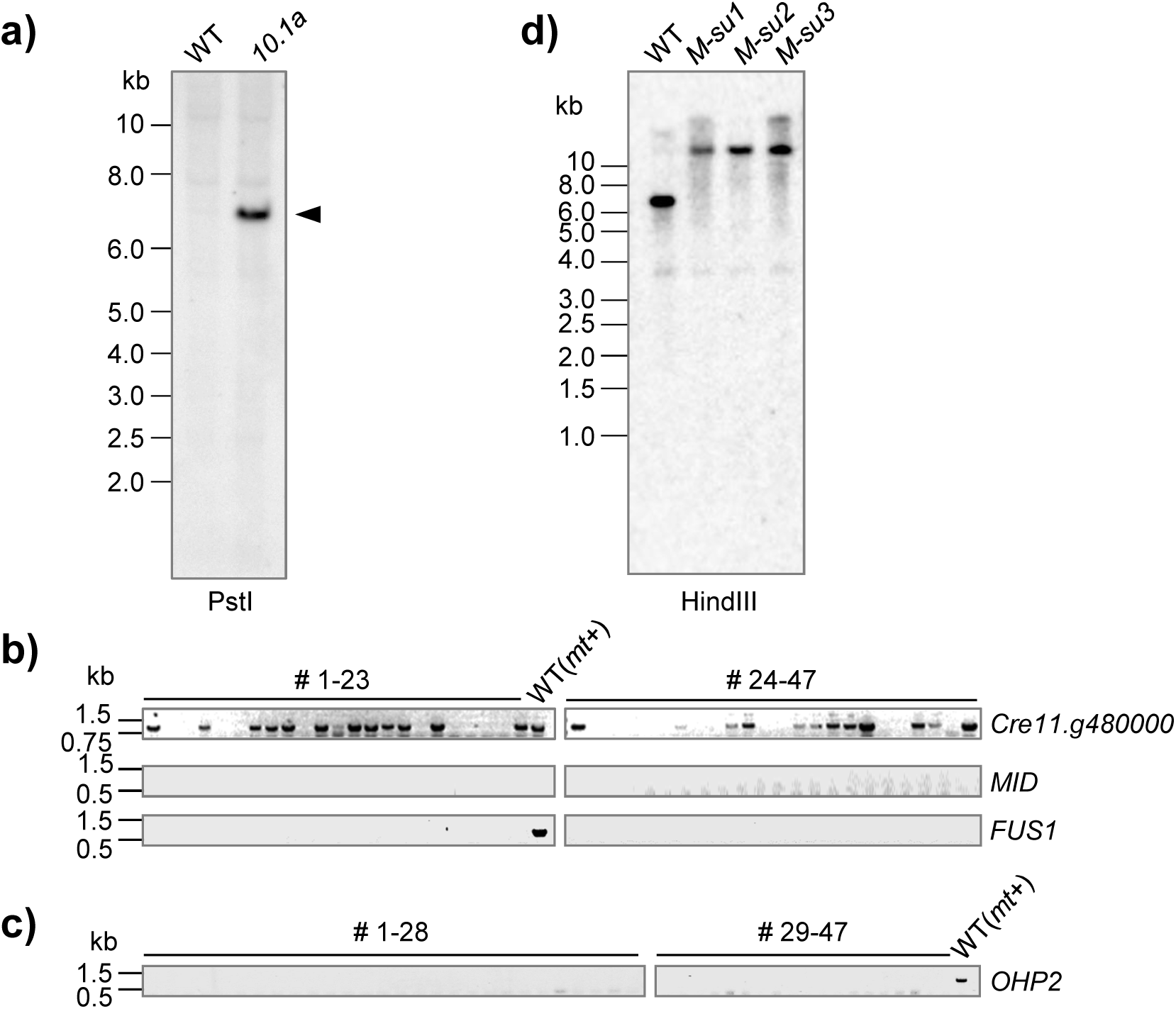
Genetic analysis of the *10.1a* mutant and suppressor strains. **a) Southern blot analysis indicates a single insertion of the mutagenic vector pBC1 in the *10.1a* mutant.** 10 µg of genomic DNA of the wild type Jex4 (WT) and the mutant strain *10.1a* were fractionated in an 0.8% agarose gel after digestion with PstI. The gel was blotted onto a nylon membrane, and hybridized with a dig-labelled probe specifically detecting the *aphVIII* gene. Molecular size markers are labeled at the left (M). Primers used to generate the probe were pBC1-APHV-fw / pBC1-APHV-rev. **b), c) Genotyping of 47 non-photosynthetic clones resulting from a backcross of the *10.1a* mutant to the WT strain WT-S34 mt*+*. b)** Analysis of PCR products using primers specific for the locus *Cre11.g480000* and mating type specific loci *MID* and *FUS1*. Upper panel: A *Cre11.g480000*-specific product of 829 bp is expected for the wild type. For clones possessing the insertion of the Pm-resistance cassette no product is expected due to the loss of the primer binding site. Only approximately half of the PSII deficient clones tested exhibited the identified insertion of the resistance cassette. The middle and lower panels show the detection of mating type specific *MID (mt*-) and *FUS1* (*mt*+) genes. Primers applied for PCR reactions were as follows: *Cre11.g480000* (hit2fw / hit2rev); *MID* and *FUS1* genes (mid-fw/mid-rev; fus1-fw/fus1-rev). Molecular size markers are indicated on the left. **c)** Analysis of PCR products using primers specific for the *OHP2* gene (*Cre06.g251150*). A product of the expected size could only be detected in the WT but not in the 47 non-photosynthetic clones investigated in b). Position of OHP1-fw2 (fw)/OHP1-rev2 (rev) primers used is indicated in Figure 1a. **d) Southern blot analysis of genomic DNA from the Jex4 recipient strain (WT) and three independent *ohp2* suppressor mutants (*M-su1 to M-su3*)**. 10 µg of genomic DNA were fractionated in a 0.8% agarose gel after digestion by HindIII, blotted onto a nylon membrane, and hybridized with the *OHP2*-specific probe indicated in Figure 1a. The signal pattern of all suppressor mutants analyzed, resembles that of the *ohp2* mutant indicating that they still harbor the large insertion of the putative *TOC1* transposon (compare strain *10.1a* in Figure 1b). For primer sequences see Supplemental Table S2.

**Figure S2.**
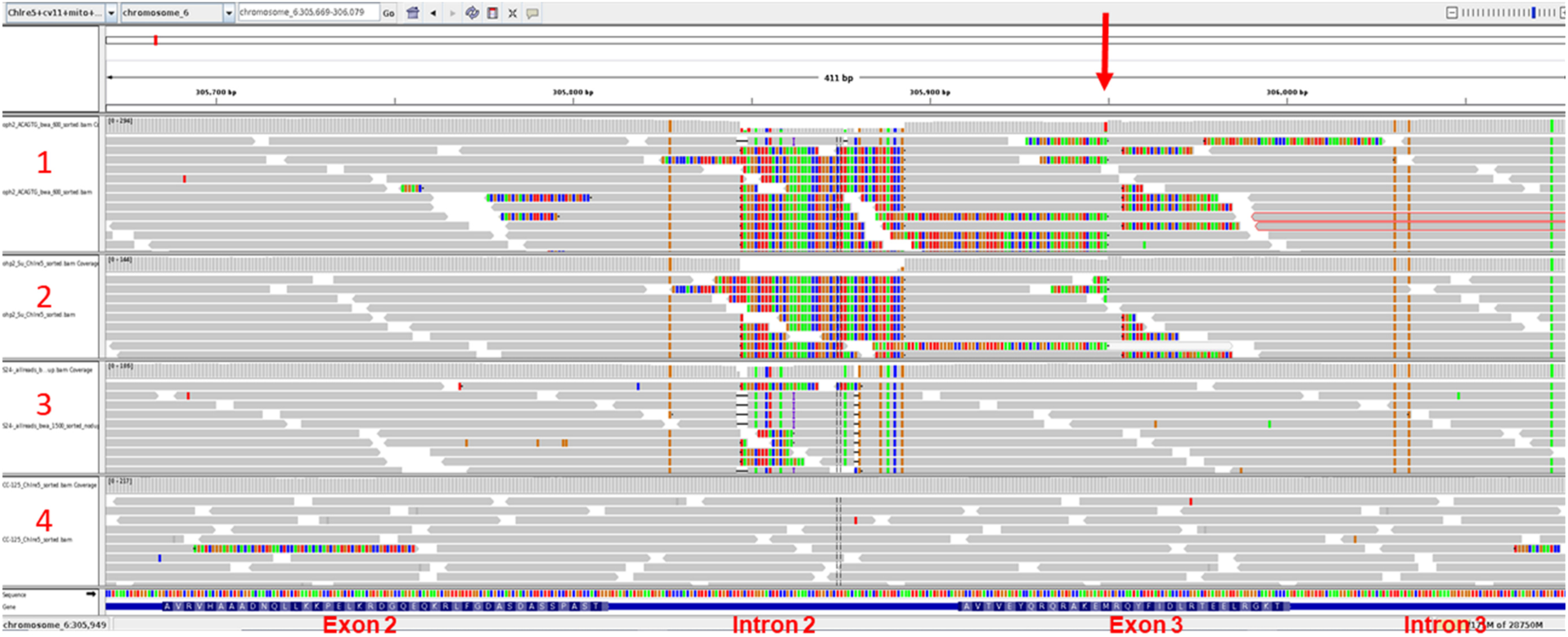
Identification of the *ohp2* mutation. Snapshot from the IGV browser showing the second and third exons of *OHP2* with mapping of genomic Illumina reads from (top to bottom) *ohp2* (1), a suppressed *ohp2* mutant (2) and the 137c WT strains WT S24-(CC-5100, Gallaher et al. 2015) (3) and CC-125 (4). The gene structure ranging from exon 2 to intron 3 is indicated in the bottom panel. Note presence of multiple polymorphisms common to all the strains from the Paris lab (top 3), including a major variant in intron 2. The *TOC1* insertion in *OHP2* is indicated by a red arrow.

**Figure S3.**
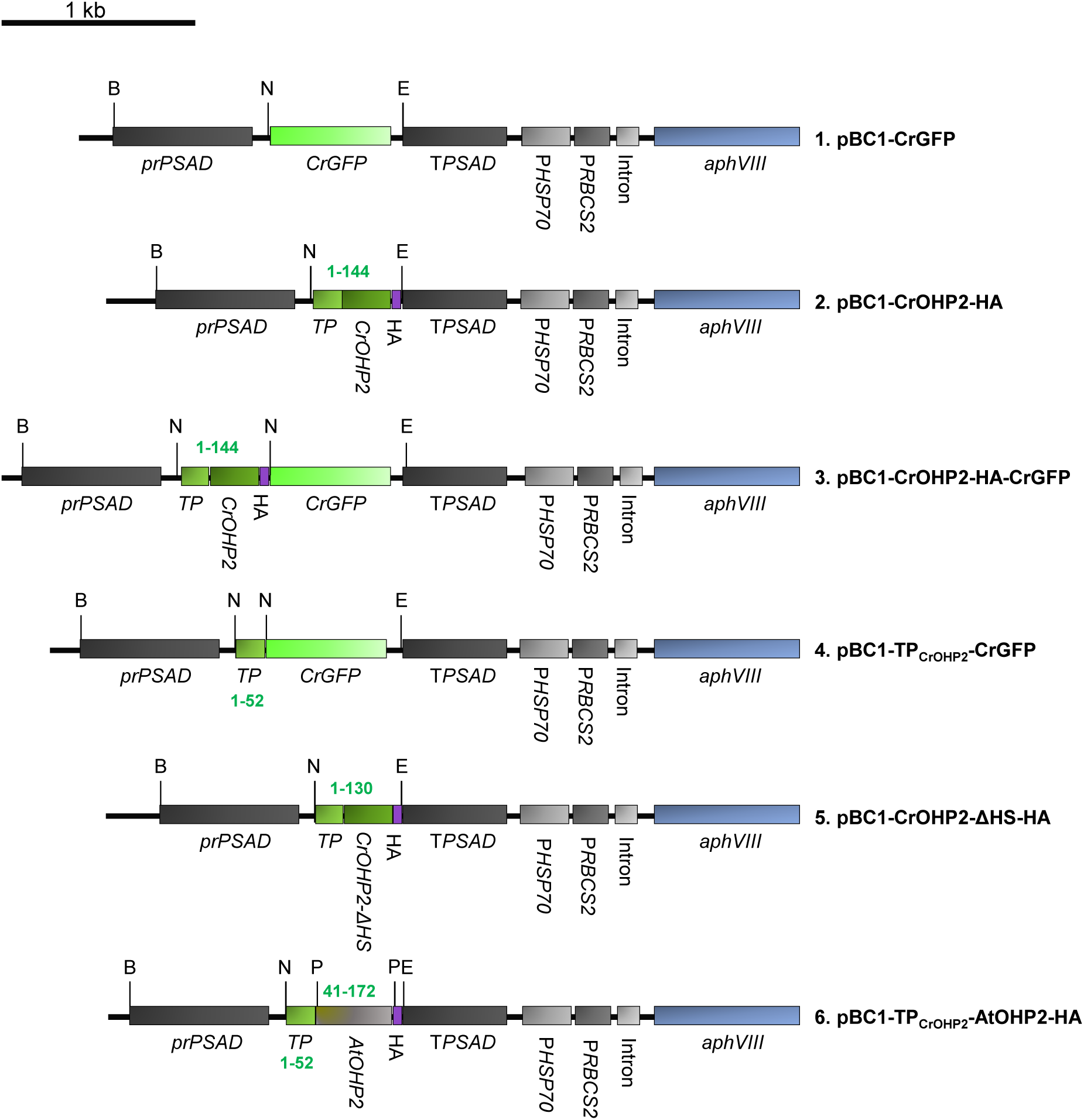
DNA constructs used for complementation and localization studies. Illustrations show the expression cassette of the pBC1-CrGFP vector used for cloning in 5’ → 3’ direction (= pJR38, Neupert et al. 2009). Sequences of interest were inserted downstream of the strong nuclear *PSAD* promoter (pr*PSAD*) via NdeI or NdeI/EcoRI restriction sites. Pm-resistance “*aphVIII*” is under the control of a fusion promoter consisting of the promoter of the gene encoding the small subunit of Rubisco (P*RbcS2*) and the gene encoding the heat shock protein 70A (P*HSP70A*). *CrOHP2* was amplified from cDNA generated from the WT strain Jex4. All constructs contain the sequence encoding the CrOHP2 transit peptide (TP) predicted by TargetP1.1 (Emanuelsson et al., 2007) to guarantee proper localization of the expressed proteins. Numbers given in green represent amino acid ranges encoded by the respective DNA insert. Important restriction sites are indicated (B: BamHI, N: NdeI, E: EcoRI, P: PstI). Primers used for cloning are given in Supplemental Table S2. For cloning details of the synthetic *Arabidopsis* gene in construct 6, see Supplemental Data S2.

**Figure S4.**
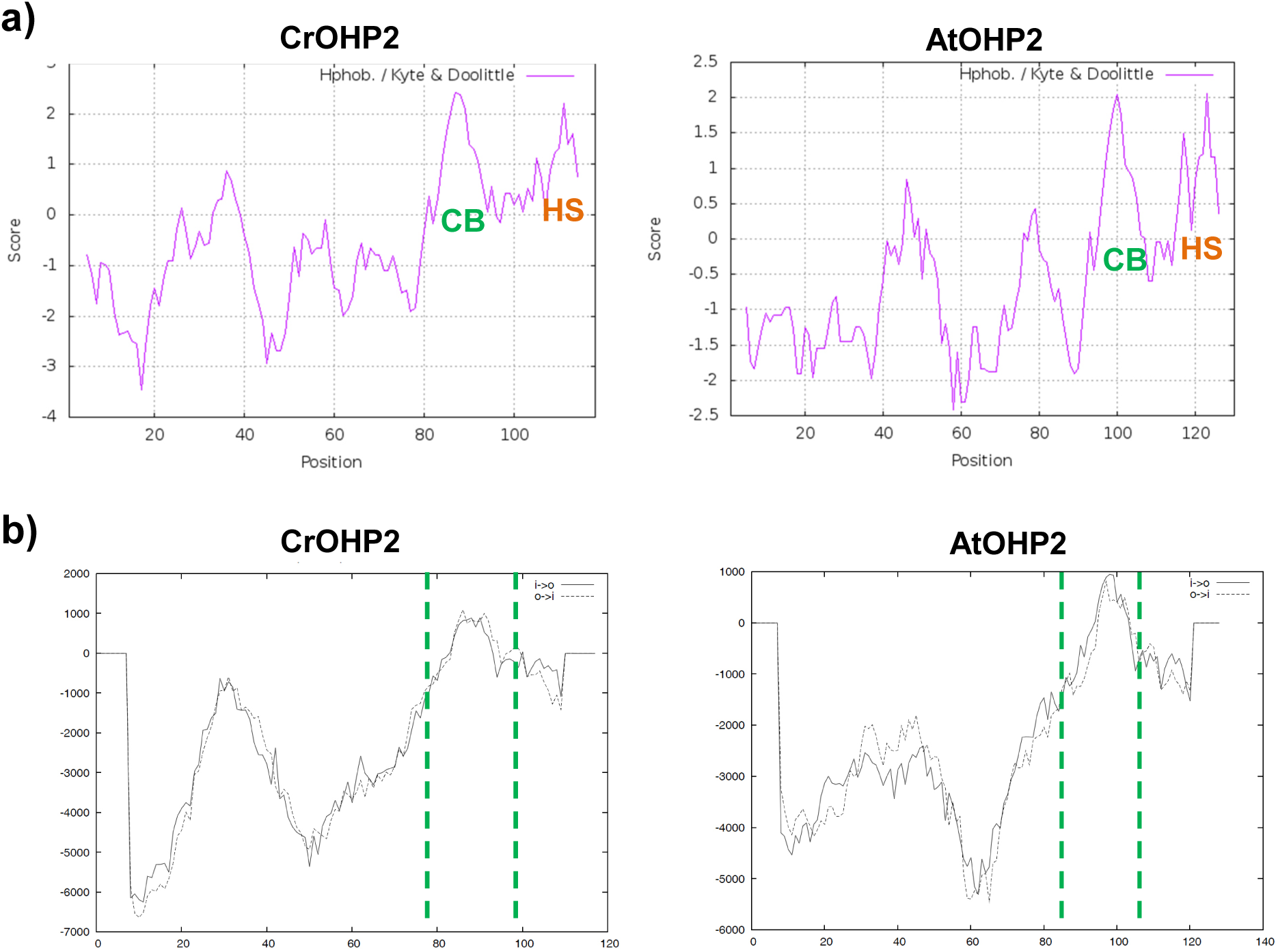
Hydrophobicity prediction for *Chlamydomonas* and *Arabidopsis* OHP2 amino acid sequences. Predictions were performed for proteins lacking the transit peptides predicted by TargetP 2.0 (*C. reinhardtii*: CrOHP2 aa 27-144; *A. thaliana*: AtOHP2 aa 44-172) **a) Hydrophobicity plot.** Indicated is the C-terminal chlorophyll-binding region (CB) and a hydrophobic stretch (HS). Hydropathy was predicted using the *ProtSacle* online software (https://web.expasy.org/protscale/) based on the model from Kyte et al. (1982). X-axis: position of amino acids; Y-axis: hydrophobicity value. **b) Prediction of membrane spanning segments**. The region between the green dashed lines is predicted to represent membrane-spanning regions (TMPred; Hofmann and Stoffel 1993; https://embnet.vital-it.ch/software/TMPRED_form.html). The X-axis represents the position of amino acids, the ordinate the TMPred score. Positive scores suggest hydrophobic regions.

**Figure S5.**
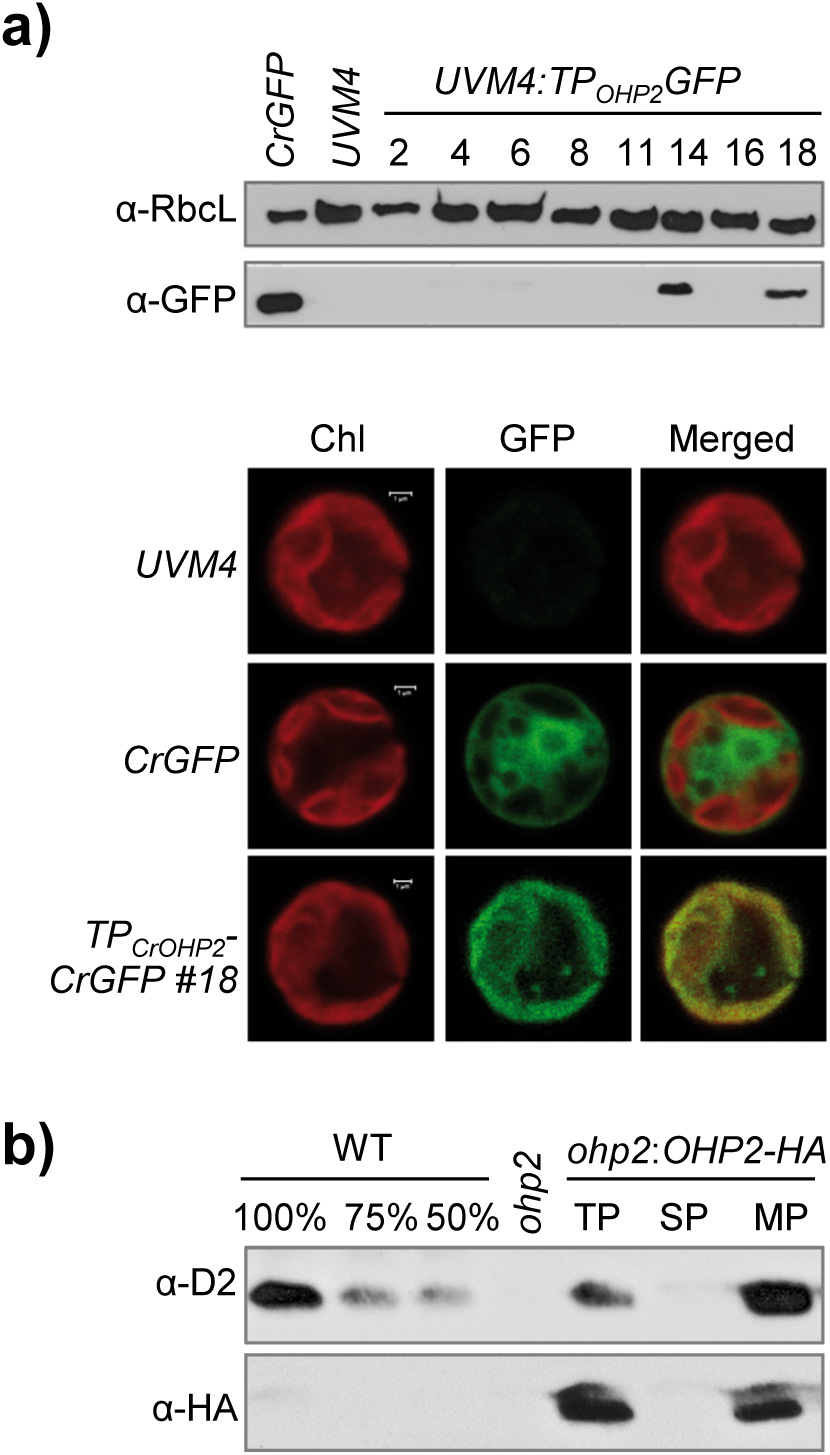
OHP2 is a membrane localized chloroplast protein. **a) GFP localization studies.** Upper Panel: Immunoblot analysis of *UVM4* strains transformed with pBC1-TP_CrOHP2_-CrGFP (construct 4, Supplementary Figure S3). 20 µg of total proteins of eight Pm-selected transformants (*UVM4*:*TP_OHP2_-GFP*) were analyzed for GFP accumulation (α-GFP) along with the untransformed recipient strain *UVM4* as negative control. 5 µg total protein of pBC1-CrGFP (construct 1, Supplementary Figure S3) transformed *UVM4* cells previously shown to accumulate GFP in the cytosol (*CrGFP*) served as positive control (Jalal et al., 2015). The protein detected in *UVM4*:*TP_OHP2_-GFP* transformants migrated slightly slower than the signal detected in the strain expressing the unfused 27 KDa cytosolic CrGFP, suggesting partial retention of the fused transit peptide. RbcL was used as loading control (α-RbcL). Lower panel: The GFP expressing clone #18 was used for GFP localization studies (*TP_OHP2_-CrGFP, clone #*18) by laser scanning confocal fluorescence microscopy using a TCS-SP5 system (Leica). The recipient strain *UVM4* as well as *CrGFP* served as controls. Chlorophyll auto fluorescence (Chl), GFP fluorescence (GFP), and the merged signals of *Chlamydomonas* cells are shown. Bar = 1 µm. **b) Cell subfractionation.** Immunoblot analysis of cell fractions from the complemented strain *ohp2:OHP2-HA* clone #53 with a D2 (upper panel) or an HA specific antibody (bottom panel). 10 µg of total (TP) and fractionated into soluble (SP) and membrane (MP) proteins of the complemented strain, as well as *ohp2* as negative control were analyzed. A dilution series of the wild type Jex4 (WT) served as reference.

**Figure S6.**
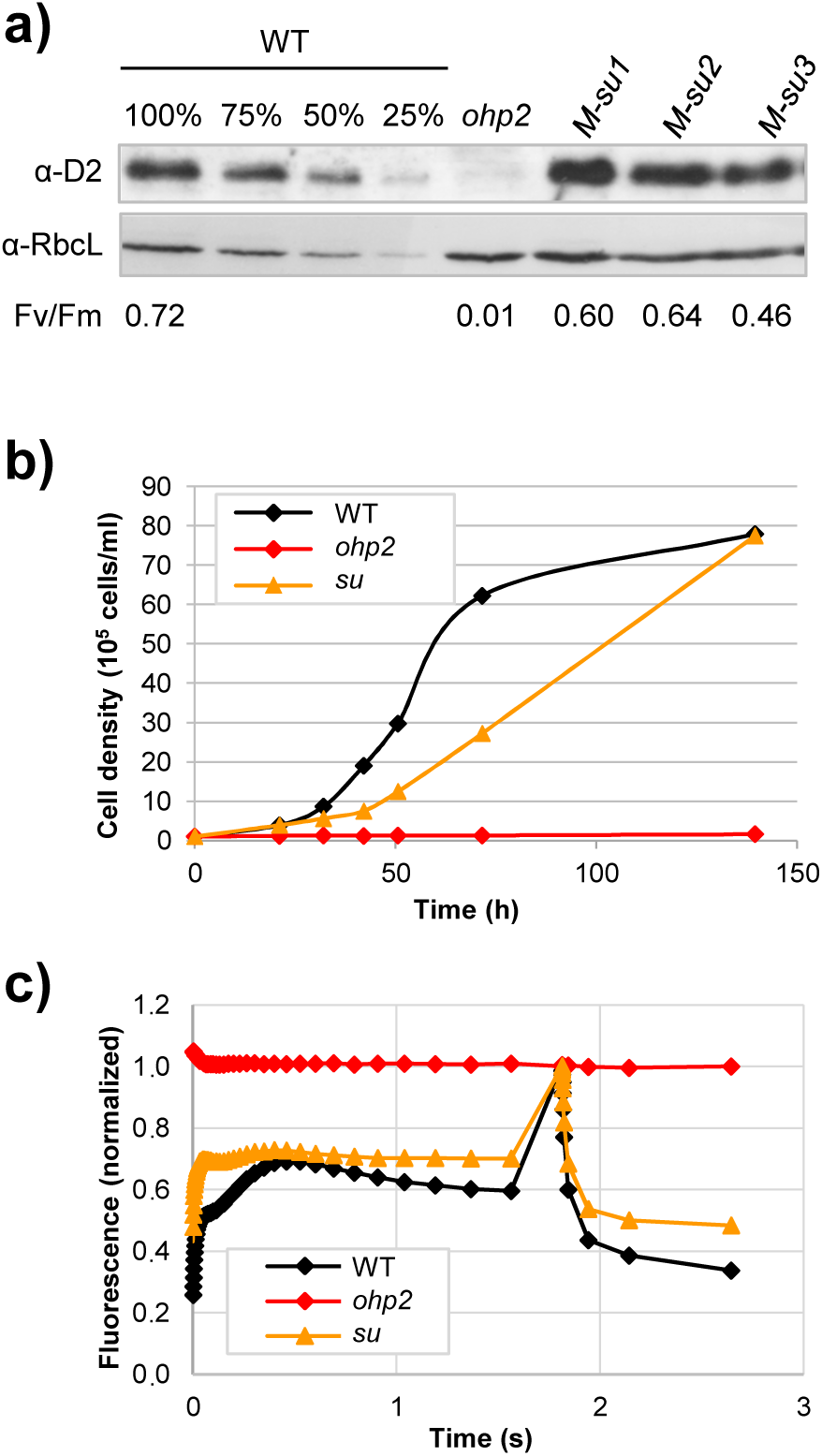
Re-accumulation of D2 and restoration of photoautotrophy in suppressor mutants. **a) Western blot.** 20 µg of whole cell proteins of three suppressor mutants (*M-su1-3*), as well as the *ohp2* strain and a dilution series of the wild type Jex4 (WT) were immunodecorated with a D2 specific antibody. RbcL served as a loading control. To measure Fv/Fm ratios given below the panel, cells were resuspended in H_2_O up to a concentration of 10^5^ cell/mL and 10 µL were spotted onto TAP plates and grown for 7d under low light at 30 µE/m^2^/s. **b) Growth curves under photoautotrophic conditions at 60 µE/m^2^/s** and **c) fluorescence induction kinetics** measured under illumination at 135 µE/m^2^/s of the WT, *ohp2*, and a suppressor strain (*su*).

**Figure S7.**
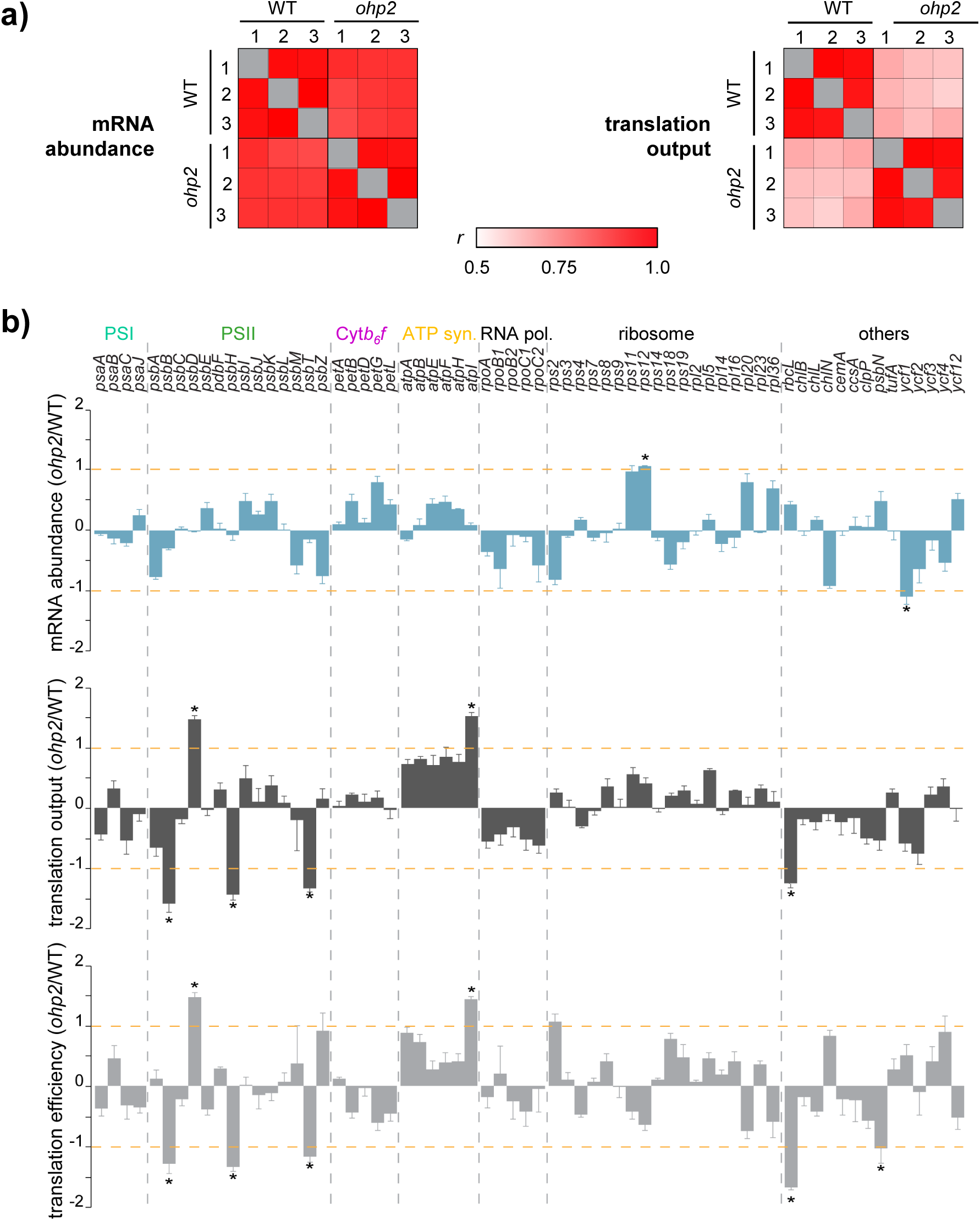
Reproducibility of RF experiments and results for all chloroplast encoding genes. **a) Reproducibility of ribosome profiling experiments.** Heat map representing Pearsońs *r* correlation values. Correlations are based on mRNA abundances and on translation output for which all probes were averaged for each Open Reading Frame, respectively. Color code is shown on the bottom. For the individual *r*-values, see Supplemental Dataset 1. **b) Average transcript abundance, translation output and translation efficiency ratios between WT and the *ohp2* strain for all chloroplast protein encoding genes, grouped by function.** The relative average mRNA abundances (upper panel), translation output (middle panel) and translation efficiency (lower panel) were calculated for each chloroplast reading frame in both *Chlamydomonas* strains, normalized to overall signal intensities (for details see Methods), and plotted as ratios (*ohp2* over WT) in log2 scale. Error bars represent standard deviations of n= 3 biologically independent samples (note that for better visualization error bars are only shown unidirectional). Asterisks indicate significant changes for genes whose RNA, protein synthesis or translational efficiency levels changed more than two-fold in the mutant, with *q*-values <0.05 (see Methods). The threshold is indicated by a horizontal orange line. Gene groups are PSI and PSII, the Cyt*b*6*f* complex, the plastid-encoded RNA polymerase (RNA pol.) and chloroplast-encoded ribosomal proteins (ribosome).

**Figure S8.**
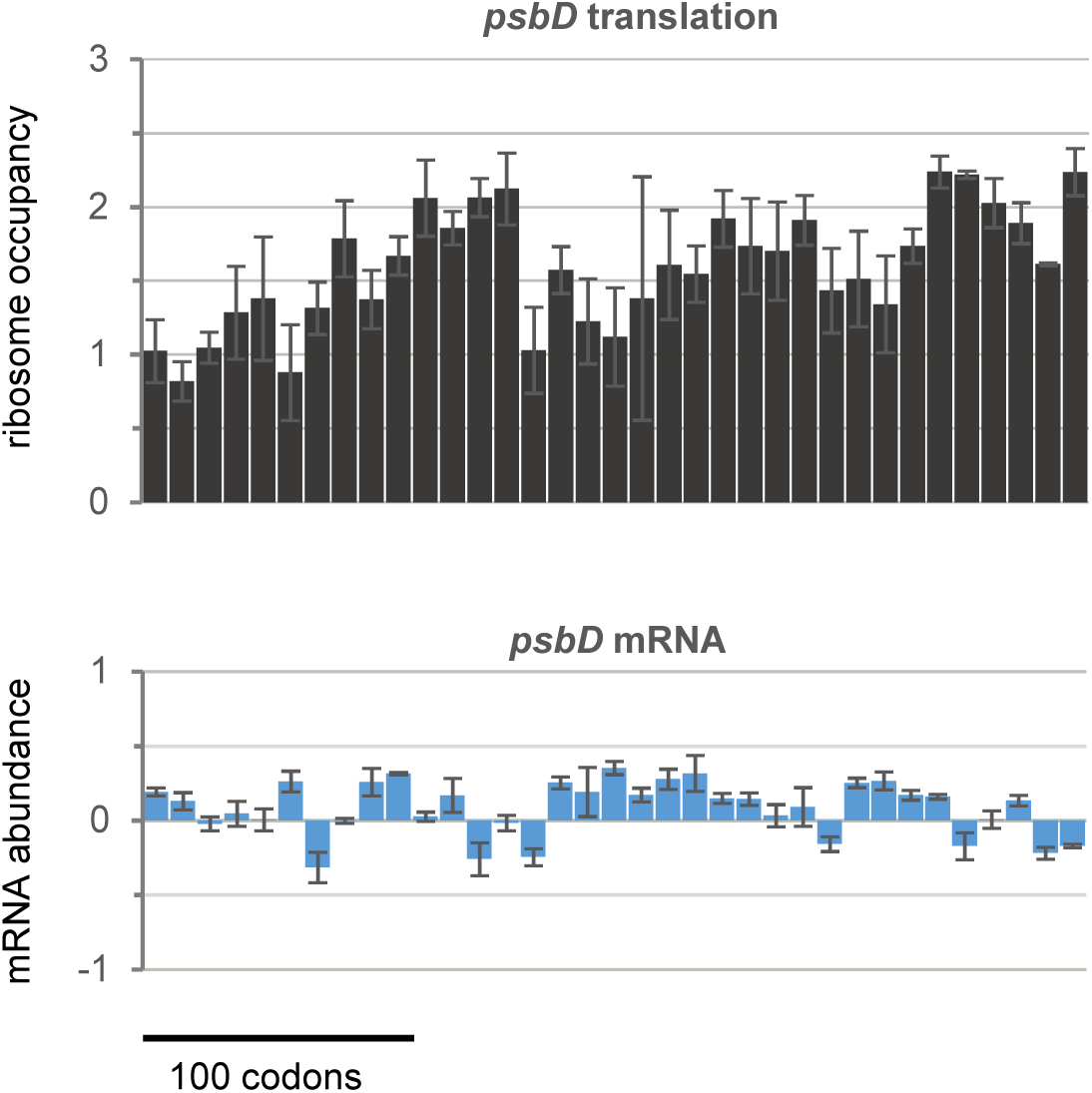
Targeted ribosome profiling of chloroplast translation reveals enhanced *psbD* translation in the *ohp2* mutant. For the Open Reading Frame of *psbD*, normalized ribosome footprint intensities were plotted as mean log_2_ ratios between *ohp2* mutant and wild type. Error bars denote differences between three independent biological replicates.

**Figure S9.**
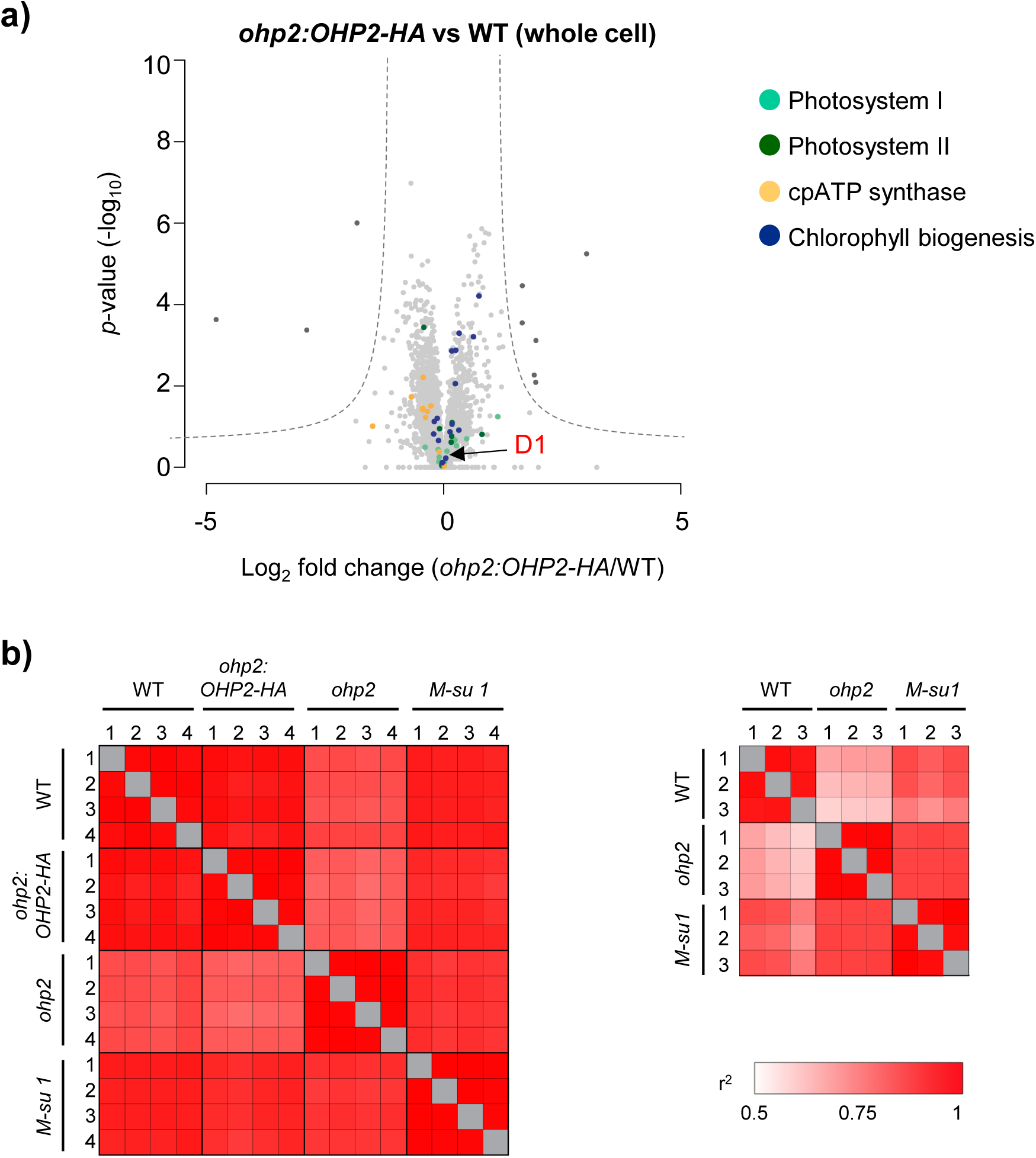
Complementation of *ohp2* and heat maps representing the reproducibility of LC-MS experiments. **a) Complementation of *ohp2* with HA-tagged OHP2 fully reverts photosynthesis-deficient phenotype.** Volcano plots represent the relative whole cell proteome changes comparing lysates of *ohp2:OHP2-HA* versus wild type strains. All experiments were performed in four independent biological replicates. Mean fold change of LFQ values (in log_2_) is plotted on the x-axis, *p*-values (in −log_10_) are plotted on the y-axis. Light grey dots represent proteins with no significant change, dark grey dots show proteins that are significantly different with FDR <0.05 and S_0_=1. Proteins of the PSI, PSII, the chloroplast ATP synthase and of proteins involved in chlorophyll biogenesis are marked in color. Large colored dots are significantly different. **b) Heat maps representing correlations between experiments of LC-MS of whole cell (left) and membrane fractions (right).** Correlations are given as r-squared LFQ values (Supplemental Datasets 2, 3). Proteins were identified through LC-MS as described in Methods.

**Figure S10.**
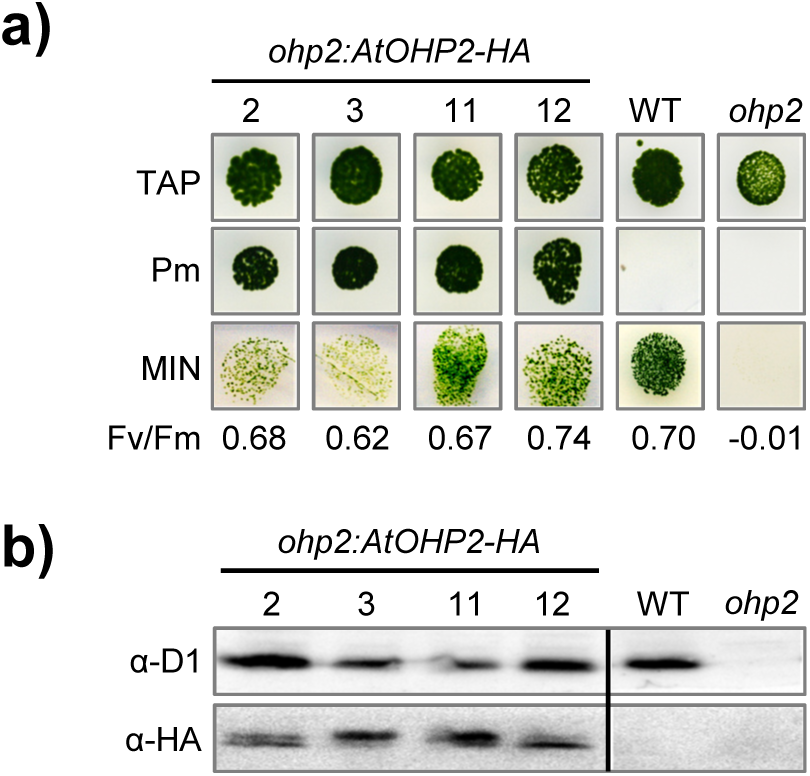
Complementation of the *ohp2* mutant with the orthologous *Arabidopsis* protein restores photoautotrophy and D1 accumulation. **a) Growth test.** The *ohp2* mutant was transformed with the construct pBC1-TP_CrOHP2_-AtOHP2-HA (see construct 6 in Supplemental Figure S3). Transformants were first selected on paromomycin (Pm) and then tested for photoautotrophic growth on HSM plates (MIN). Four representative clones out of 20 are shown. Fv/Fm values are indicated below the panel. **b) Immunoblot analysis** of Pm-resistant transformants. 20 µg of total proteins were separated on 15% denaturing polyacrylamide gels and analyzed with the antibodies indicated on the left. Jex4 (WT) and *ohp2* were run on the same gel but not in adjacent lanes as indicated by a vertical black line.

**Figure S11.**
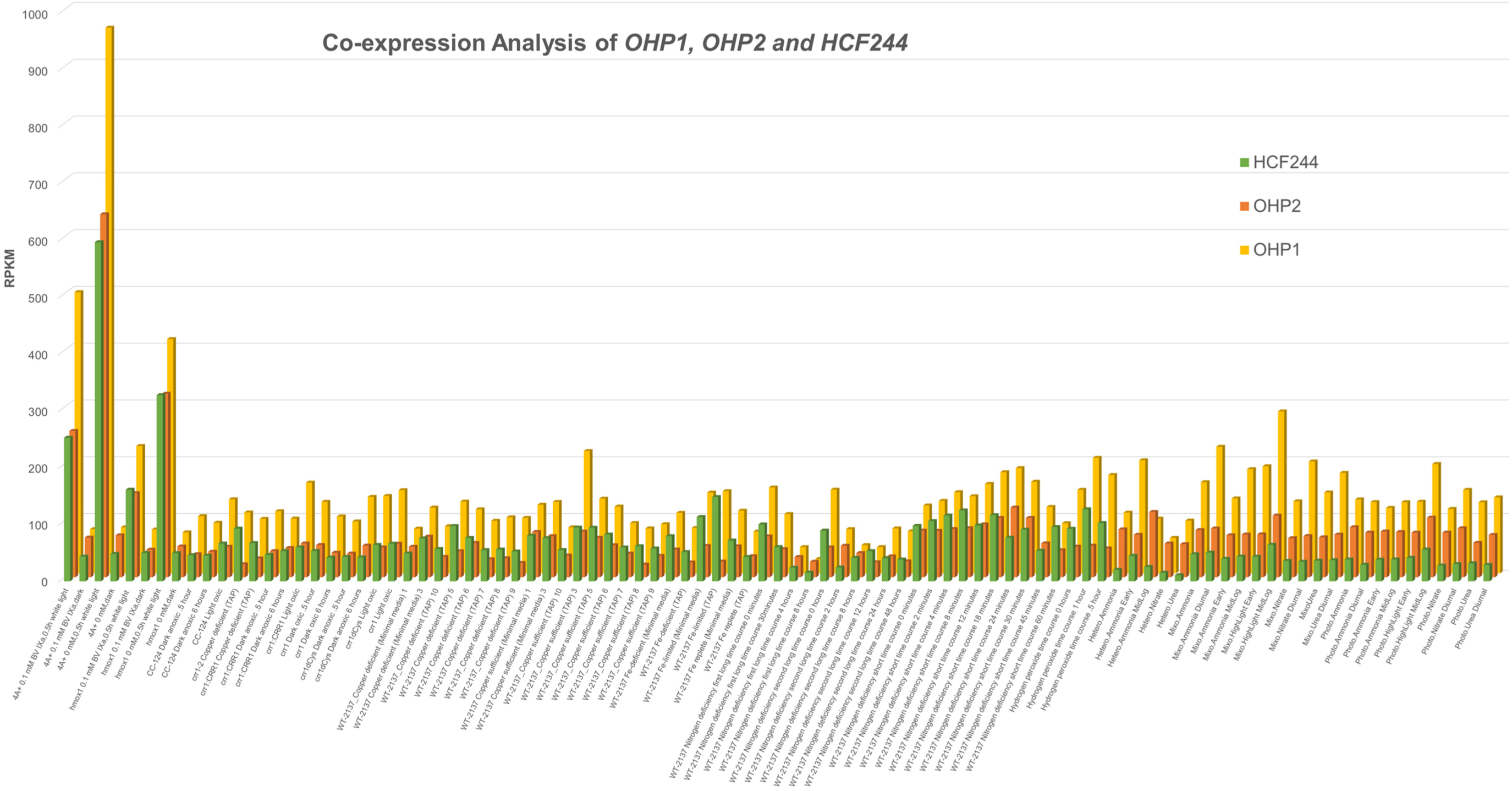
Transcript accumulation level of *OHP2* (*Cre06.g251150*), *OHP1* (*Cre02.g109950*), and HCF244 (*Cre02.g142146*) in *Chlamydomonas* under various growth conditions. Transcriptomic data originate from independent studies (Hemschemeier et al. 2013; Duanmu et al. 2013; Castruita et al. 2011; Urzica et al. 2012a,b; Boyle et al. 2012) and were obtained from the InterMine interface on Phytozome. Mean values are expressed in reads per kilobase per million (RPKM).

### SUPPLEMENTAL TABLES, DATA AND REFERENCES

**Supplemental Table S1.** Photosynthetic parameters and Chl composition of *Chlamydomonas* WT Jex4 (WT), the *ohp2* mutant and *ohp2* complemented with OHP2-HA (construct 2, Supplemental Figure S3; strain *ohp2:OHP2-HA*, clone #53). Photosynthetic parameters were determined as described in Experimental Procedures. Chl determination was performed according to Arnon (1949).

| Strain | WT | *ohp2* | *ohp2:OHP2-HA* |
| --- | --- | --- | --- |
| **Photosynthetic parameters:** |  |  |  |
| Fluorescence: Fv/Fm | 0.77 | -0.04 | 0.79 |
| ECS: PSII/PSI | 1.40 | 0.05 | 1.50 |
| ECS: phase b/phase a | 0.40 | 0.47 | 0.50 |
| **Chl determination (n= 3):** |  |  |  |
| Chl *a* (pg/cell) | 4.17 ± 0.12 | 2.31 ± 0.11 (55%) | 3.32 ± 0.27 (80%) |
| Chl *b* (pg/cell) | 2.25 ± 0.12 | 1.44 ± 0.08 (65%) | 1.70 ± 0.10 (75%) |
| Chl *a + b* (pg/cell) | 6.65 ± 0.24 | 3.75 ± 0.19 (58%) | 5.00 ± 0.37 (78%) |
| Chl *a/b* | 1.85 | 1.61 | 1.95 |

**Supplemental Table S2:**
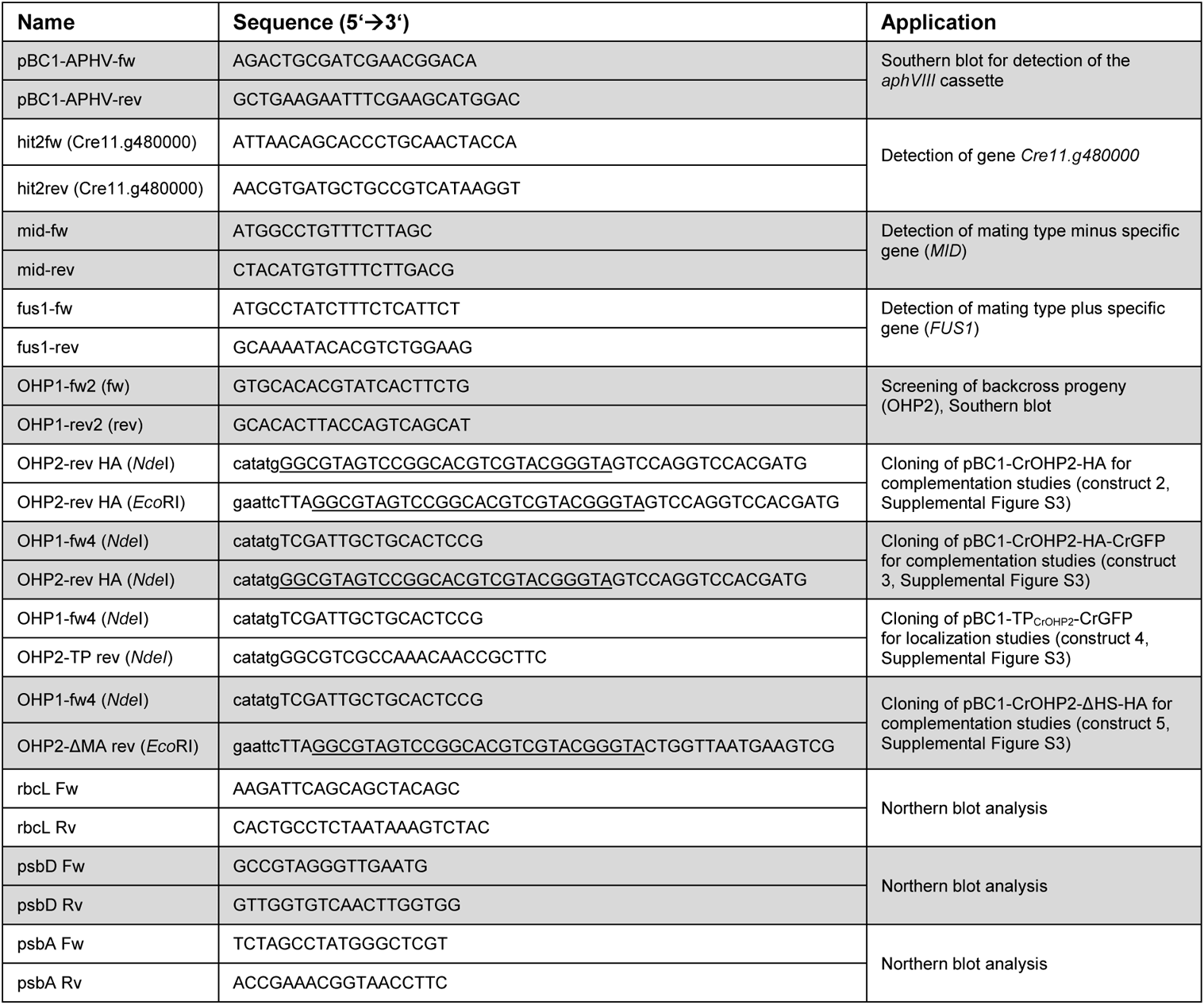
Primers used in this study. Restriction sites in lower case letters. Sequence coding for an HA-tag is underlined.

**Supplemental Data S1. Flanking sequence tags in the *ohp2* mutant obtained by Illumina sequencing.** The *ohp2* mutant shows a putative TOC1 transposon insertion in the third exon of the *OHP2* gene between position chromosome_6:305949 and 305950. Nucleotides shown in blue correspond to genome sequences (left side: chromosome_6:305923..305949; right side: 305950..305975) while those in red are identical to LTR sequences described for *TOC1* retrotransposons.

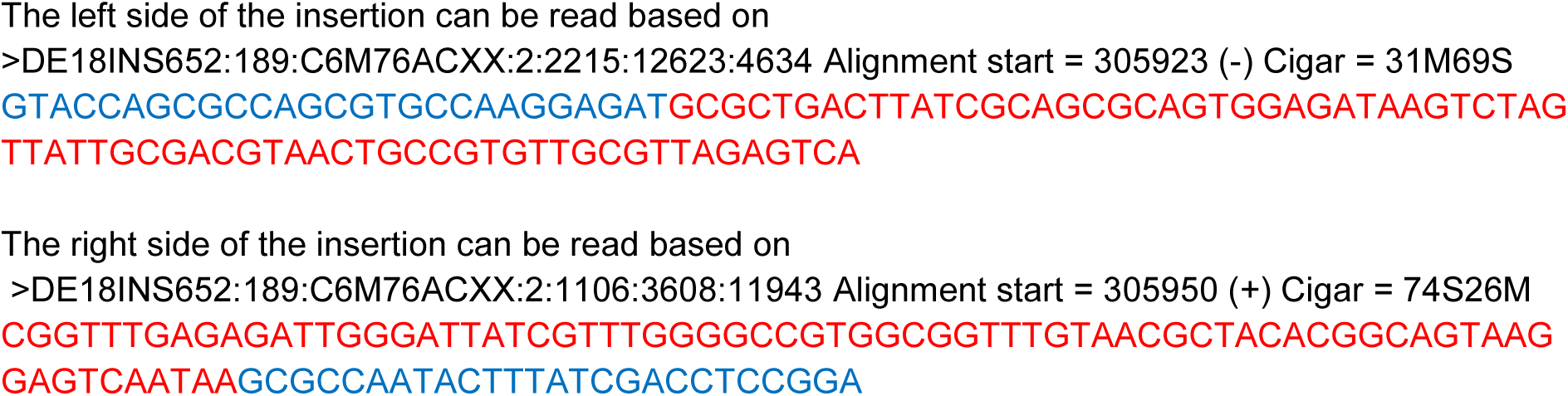

**Supplemental Data S2. Synthetic Arabidopsis** *OHP2* **nucleotide sequence and derived protein sequence used to complement the** *Chlamydomonas ohp2* **mutant strain.**

a) Sequence of codon adapted synthetic gene from *Arabidopsis* ordered from Integrated DNA Technologies (IDT) that was used to clone the construct pBC1-TP_CrOHP2_-AtOHP2-HA (construct 6 in Supplemental Figure S3). In green the *Chlamydomonas* sequence encoding the N-terminal region including the transit peptide (aa 1-52, *Cre06.g251150*) predicted by TargetP-1.0. In cyan, the sequence encoding the *Arabidopsis* protein OHP2 (aa 41-172, *AT1G34000*) adapted to the codon usage of *Chlamydomonas*. The HA-tag encoding sequence is shown in purple letters. Restriction sites introduced for cloning are displayed in lower case letters.

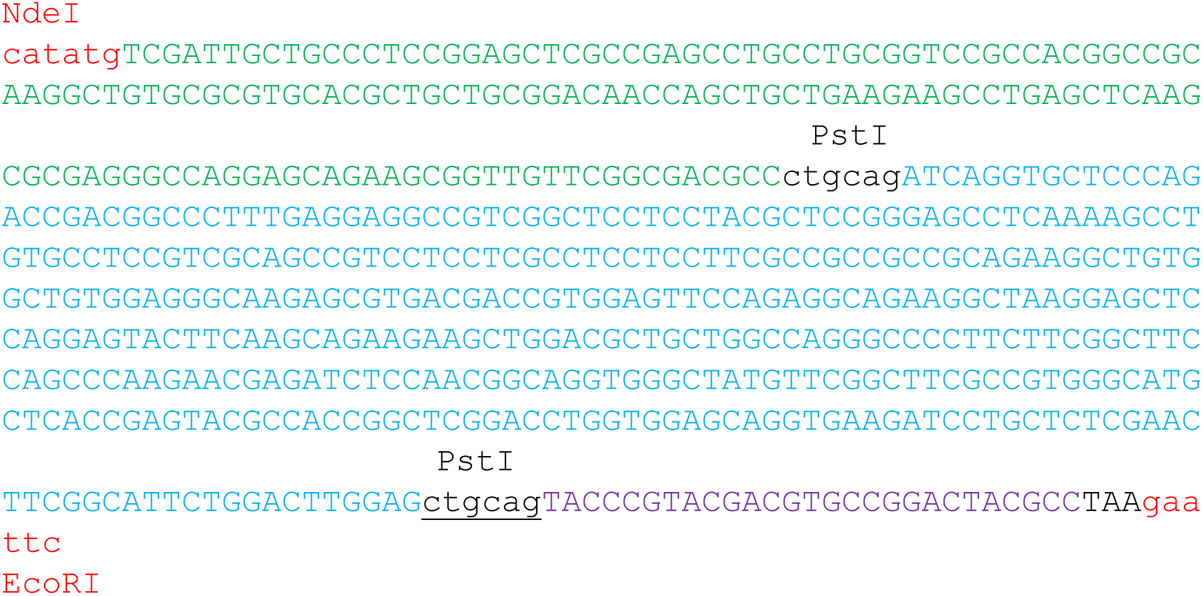

b) Chimeric protein sequence expressed from *Chlamydomonas* strains transformed with the vector pBC1-TP_CrOHP2_-AtOHP2-HA. Color code as in a). The fusion protein possesses the *Chlamydomonas* N-terminus fused to the C-terminally HA-tagged *Arabidopsis* protein lacking the predicted localization signal.

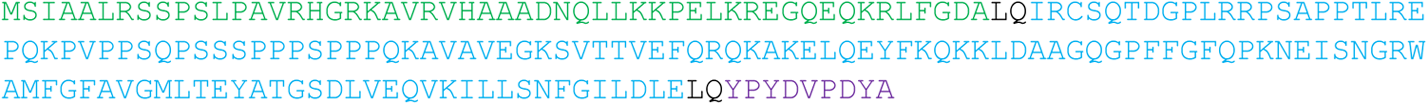

